# Predicting the synaptic partners of masked neurons across two female *Drosophila* connectomes with per-region graph models

**DOI:** 10.64898/2026.11.23.768740

**Authors:** Ahmadh Hassan, Tyler Ho, Ali Aslam

## Abstract

Whole-brain connectomes of adult *Drosophila* now exist for several individuals, so one can ask how accurately a neuron’s synaptic partners can be imputed in one animal by models trained on another. We trained a separate graph neural network for each of twelve brain regions on the FlyWire female brain and tested whether it could recover the connections of masked neurons in a second female brain reconstructed by a different pipeline (BANC). Every edge of a masked neuron was hidden, but the neuron kept features computed from its complete reconstruction (neuropil innervation, synapse-position features and total in-region synapse count) together with its cell type, transmitter and class labels. Each true partner was ranked against 100 non-partners drawn in proportion to their visible degree, and performance was measured by mean reciprocal rank (MRR). Under a decision rule fixed in the project’s analysis plan before the confirmation masks were scored, every regional model outperformed the best of three non-graph baselines: geometric-mean MRR ratios ran from 1.33 to 2.08, and model MRR from 0.36 to 0.60. Looking up each masked neuron’s partners through its matched FlyWire neuron, with no learning, reached MRR 0.18–0.48, and the regional models exceeded this lookup in all twelve regions. The comparison is conditional on inputs that carry information about the hidden edges: the passing models’ innervation and position features were computed from each neuron’s complete reconstruction, the second brain’s cell types were assigned by matching neurons to FlyWire, and only the graph models read the wiring of a masked neuron’s visible same-type neurons (a rule that only copies those neurons’ partners lost to the regional models on every confirmation instance). In an exploratory cross-region analysis, each region’s own model outscored all eleven models trained elsewhere, but this tracked how much of the target region’s cell-type vocabulary the visiting model knew, so it does not isolate region-specific wiring rules.

## 1 Introduction

Electron-microscopy connectomics has moved from partial volumes to complete adult brains. FlyWire reconstructed the 139,255 neurons of one female *Drosophila* brain from the FAFB volume [1, 2]. Companion work annotated every neuron with a hierarchy of labels and cell types and compared it with the partial hemibrain connectome of a second animal [3, 4]. That comparison reidentified most hemibrain cell types, although about a third could not be reliably reidentified, and about half of hemibrain type-to-type connections recurred in FlyWire; connections carried by more than ten synapses recurred more than 90% of the time [3]. Further connectomes followed: a female brain and nerve cord (BANC) [5], a male central nervous system [6], and comparisons of identified neuron classes across datasets [7]. With several reconstructed individuals in hand, the variability of wiring between animals can be measured rather than assumed.

The comparisons cited above report reproducibility mainly at the level of cell types and type-to-type connections. This study asks a neuron-level question. If every connection of a neuron is hidden in one animal, can a model trained on a different animal rank that neuron’s true synaptic partners above plausible non-partners? Success would mean that relations learned in the first animal, between cell identity, position, innervation and surrounding wiring, carry over to individual neuron pairs in the second. Failure would bound how far such learned relations transfer across individuals and reconstruction pipelines.

Prediction of connectivity from anatomy has a long history. Peters’ rule and its refinements test how far axon–dendrite proximity predicts synapses [8, 9]. Algorithmic reconstructions predict cortical connectivity from morphology [10–12]. Mouse functional connectomics has related connection probability to cell type and response properties [13–15]. Generative network models reproduce macroscale and small-circuit topology from spatial and homophily rules [16–18], and in *C. elegans* a stochastic developmental model reproduces connectivity shared across several adult worms [19, 20]. In machine learning, connectivity prediction is a link-prediction problem, often approached with graph neural networks [21–23], and graph neural networks have been applied within a single fly connectome to recover randomly held-out edges [24]. Link-prediction benchmarks have shown that results depend heavily on how negatives are chosen, because easy or degree-biased negatives let degree heuristics look strong [25, 26].

Our test differs from these in four ways. Models are trained on one individual and scored on another. A whole neuron’s connections are hidden at once rather than individual edges. Each true partner is ranked against non-partners drawn in proportion to their visible degree, so degree alone gains little. And the test runs separately in twelve brain regions under a decision rule fixed before the confirmation data were scored. We address two questions:

1. 1. *Cross-individual prediction.* How well do regional models trained on the FlyWire brain (F1) predict the partners of masked neurons in the BANC brain (F2)? We compare them with non-graph baselines that receive pairwise features built from the same annotations but read no F2 edges beyond visible synapse counts, and, after the registered comparison, with two rules that copy wiring directly from matched F1 neurons or from visible neurons of the same type in F2.
2. 2. *Regional specificity.* Do connection patterns learned in one region transfer to others, and can the cross-region comparison separate region-specific organization from differences in cell-type composition, neurotransmitter composition and model inputs?

Both questions are motivated by the broader hypothesis that differences in wiring organization between regions contribute to specialized brain functions. The experiments here do not test that hypothesis, identify which wiring patterns matter, or establish any causal link to function. As shown below, the evidence answers the first question in a conditional form and leaves the second open.

## 2 Results

### 2.1 A cross-individual gap-filling task

We used three adult connectomes (Table 1). F1, FlyWire FAFB v783, is a female brain [1, 3] and the only training data. F2, BANC v888, is a female brain and nerve cord from a different animal, reconstructed with a different synapse detector [5]; it supplied the development and confirmation tests. F3, the male CNS v1.0 [6], was used only for exploratory readouts. We grouped neuropils into twelve regions following the standard insect-brain nomenclature [27] (Table 2) and, for each region and dataset, built a directed graph whose edges are neuron pairs with enough synapses within that region’s neuropils (Methods). The F1 graphs span more than an order of magnitude in size, from 3,219 neurons in the central complex to 93,270 in the optic lobe (Fig. 2).

**Table 1:**
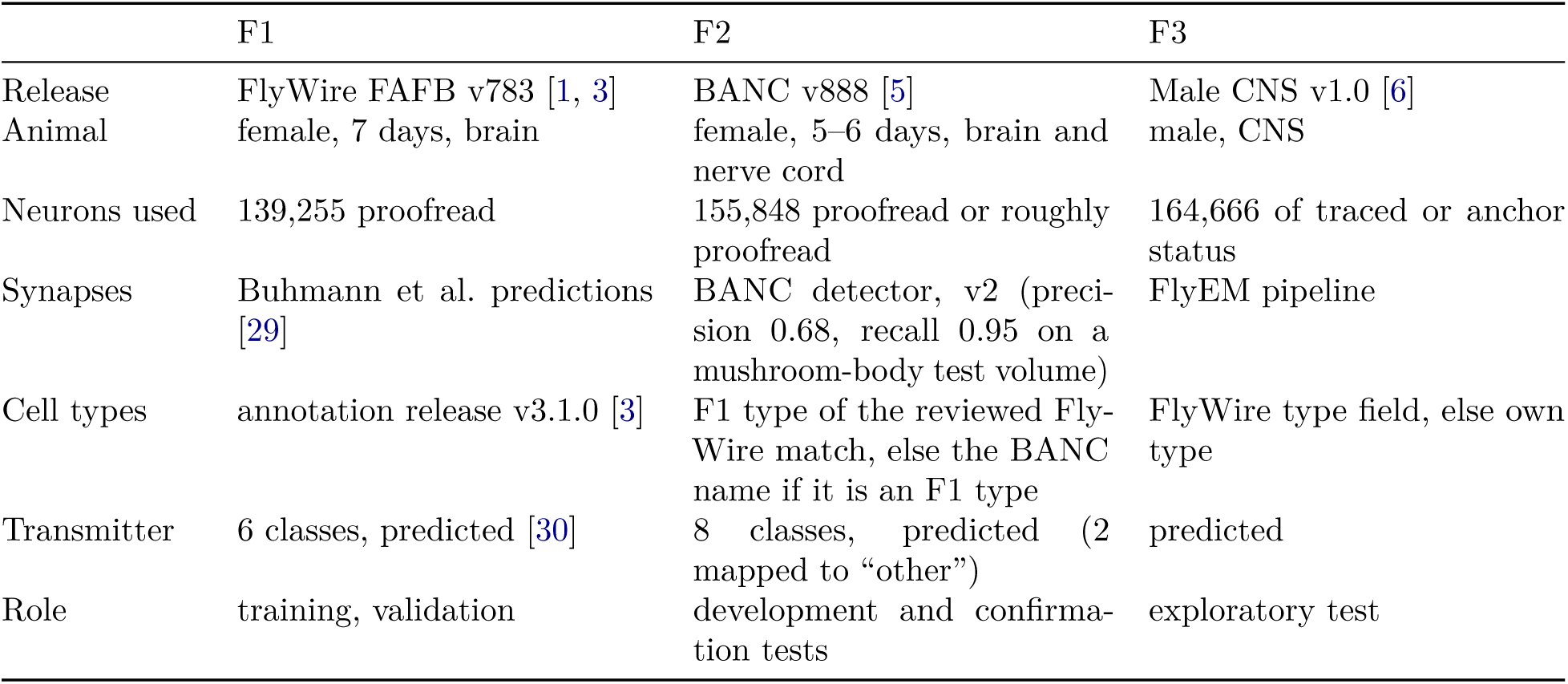
Connectomes used. Counts are for the populations this study used.

|  | F1 | F2 | F3 |
|---|---|---|---|
| Release | FlyWire FAFB v783 [1, 3] | BANC v888 [5] | Male CNS v1.0 [6] |
| Animal | female, 7 days, brain | female, 5–6 days, brain and nerve cord | male, CNS |
| Neurons used | 139,255 proofread | 155,848 proofread or roughly proofread | 164,666 of traced or anchor status |
| Synapses | Buhmann et al. predictions [29] | BANC detector, v2 (preci-sion 0.68, recall 0.95 on a mushroom-body test volume) | FlyEM pipeline |
| Cell types | annotation release v3.1.0 [3] | F1 type of the reviewed Fly-Wire match, else the BANC name if it is an F1 type | FlyWire type field, else own type |
| Transmitter | 6 classes, predicted [30] | 8 classes, predicted (2 mapped to “other”) | predicted |
| Role | training, validation | development and confirma-tion tests | exploratory test |

**Table 2:** Region graphs. An edge is a directed neuron pair whose synapse count summed over the region’s neuropils reaches the threshold: 5 in F1, and a per-region F2 threshold (Thr.) calibrated to match cell-type-pair connection densities (Methods). Types are distinct cell types among graph neurons (F2 types in F1’s vocabulary); Typed is the share of F2 graph neurons with an F1 type.

|  |  | F1 |  |  |  | F2 |  |  |  |
|---|---|---|---|---|---|---|---|---|---|
| Code | Region | Neurons | Edges | Types | Thr. | Neurons | Edges | Types | Typed |
| CX | Central complex | 3,219 | 70,830 | 488 | 3 | 3,189 | 65,514 | 476 | 99.3% |
| AL | Antennal lobe | 3,827 | 67,576 | 631 | 4 | 4,886 | 79,691 | 746 | 99.5% |
| LH | Lateral horn | 5,783 | 59,218 | 1,948 | 1 | 6,969 | 176,915 | 2,259 | 97.9% |
| MB | Mushroom body | 8,660 | 65,557 | 1,619 | 3 | 7,348 | 32,726 | 1,506 | 93.0% |
| LX | Lateral complex | 3,694 | 40,508 | 1,234 | 2 | 4,298 | 66,165 | 1,510 | 98.8% |
| SNP | Superior neuropils | 14,428 | 157,849 | 3,998 | 2 | 12,222 | 190,735 | 3,680 | 98.5% |
| INP | Inferior neuropils | 13,480 | 105,222 | 3,822 | 3 | 14,110 | 127,583 | 3,925 | 98.1% |
| VMNP | Ventromedial neuropils | 10,737 | 142,711 | 3,450 | 2 | 11,633 | 240,757 | 3,589 | 96.9% |
| VLNP | Ventrolateral neuropils | 20,179 | 361,760 | 4,572 | 2 | 20,798 | 487,419 | 4,648 | 93.6% |
| PENP | Periesophageal neuropils | 7,337 | 68,078 | 2,544 | 2 | 7,253 | 96,002 | 2,329 | 98.2% |
| GNG | Gnathal ganglia | 8,873 | 118,094 | 2,741 | 3 | 10,136 | 135,451 | 2,775 | 94.3% |
| OL | Optic lobe | 93,270 | 1,466,840 | 742 | 4 | 88,952 | 706,120 | 718 | 59.6% |

In each test instance a mask hides every edge of a set of neurons (Fig. 1). Masks come in five regimes: 1%, 5% or 10% of neurons at random (R01, R05, R10), every neuron of one cell type in both hemispheres (T, “typed”), or a spatial ball holding 5% of neurons (S05) (Fig. 3). A masked neuron keeps its cell-type label; its transmitter, superclass and class labels; the centroid and spread of its synapses in the region; the fraction of those synapses in each of the region’s neuropils (its innervation); and its total in-region synapse count, which we call the granted total. For every hidden edge with exactly one masked endpoint, the model ranks the true partner against 100 decoys. Each decoy keeps the masked neuron and replaces the partner with a non-partner of the same region graph, drawn with probability proportional to its visible synapse count on the relevant side (its in-strength if the partner is a target, its out-strength if a source). Performance is the mean reciprocal rank (MRR) of the true partners; random ranking gives 0.051.

**Figure 1:**
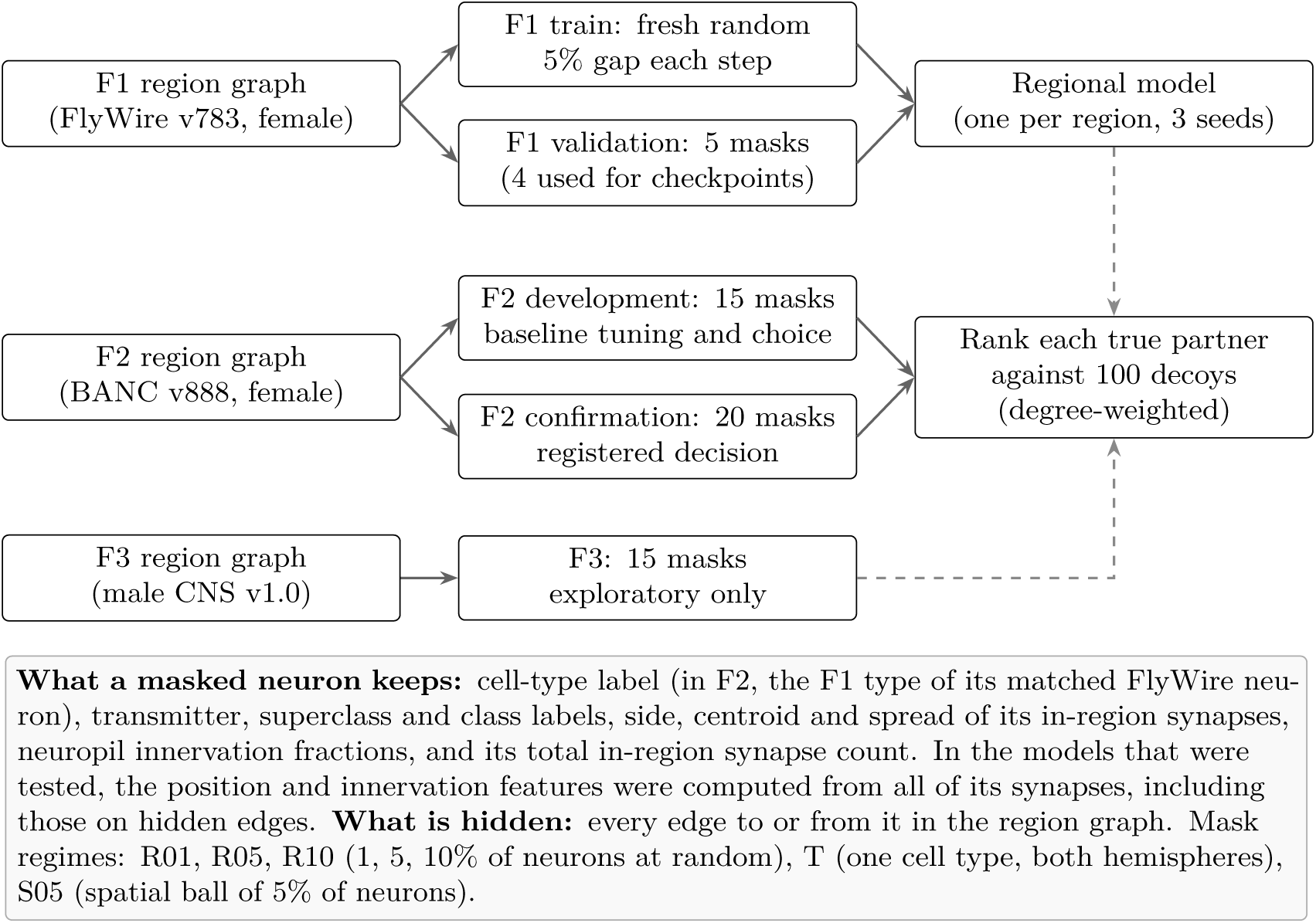
Task and data splits. Models train only on F1. F2 development masks were used to tune and choose the baselines; the registered comparison was read once per region on F2 confirmation masks. Dashed arrows mark scoring.

**Figure 2:**
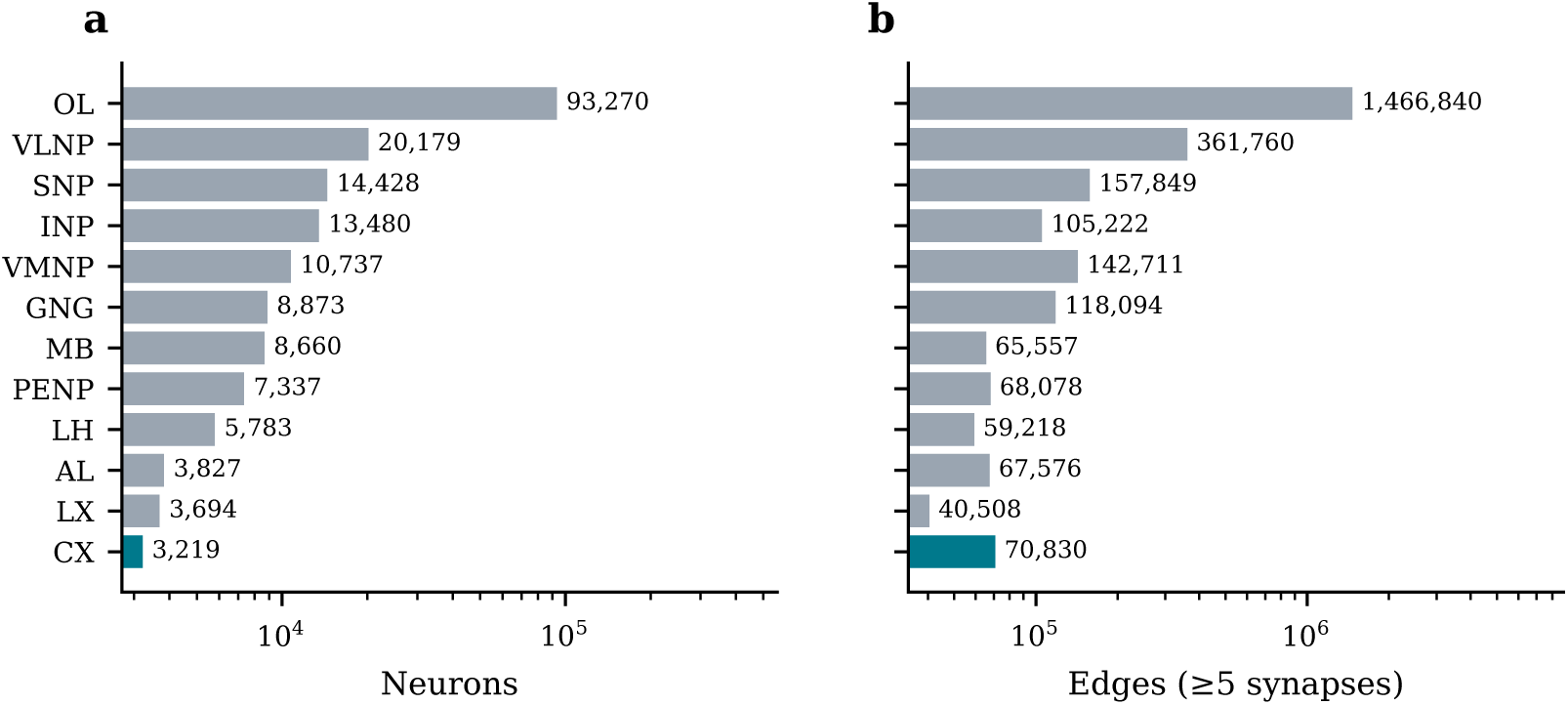
Sizes of the twelve F1 region graphs: **a**, neurons; **b**, directed edges with at least five synapses in the region (log scale). The central complex (teal) is the smallest region and was the first one studied. Counts match Table 2.

**Figure 3:**
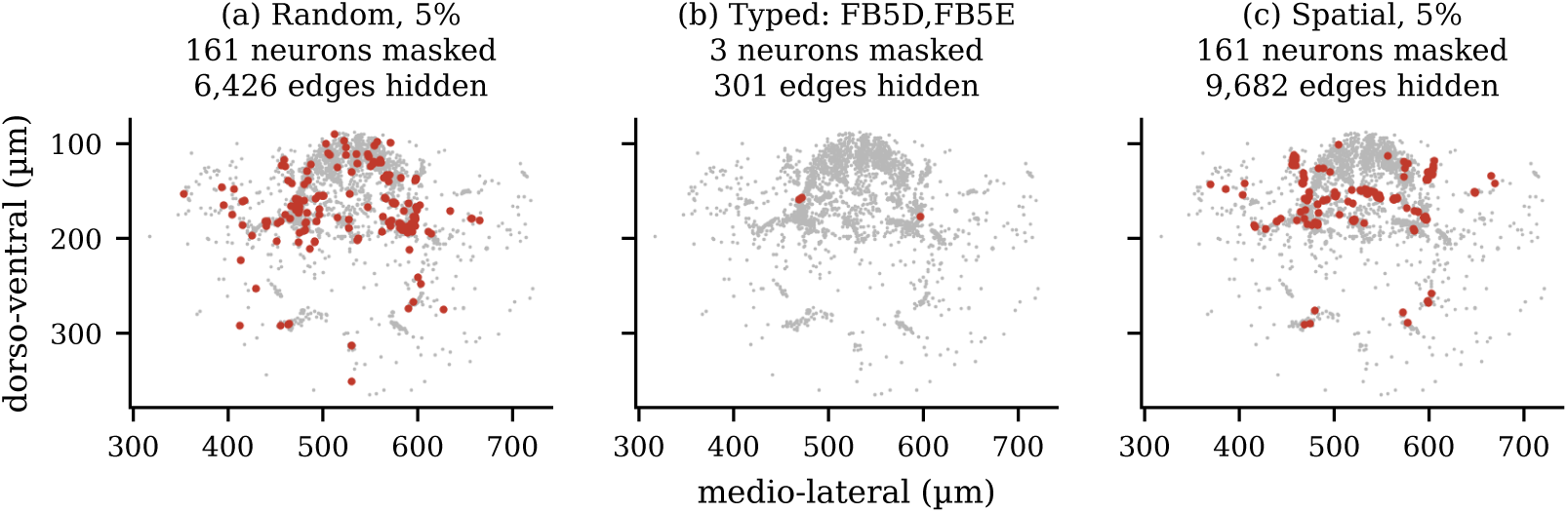
The three mask families, shown on central complex F1 validation instances. Grey points are central complex neurons at their positions (frontal projection); red points are masked neurons, all of whose region edges are hidden. **a**, random, 5% of neurons (161 neurons, 6,426 edges hidden); **b**, typed, every neuron of one cell type (3 neurons, 301 edges); **c**, spatial ball holding 5% of neurons (161 neurons, 9,682 edges). Supplementary Fig. S6 shows all five validation instances. information derived in part from the hidden wiring, and in F2 it was assigned by reference to F1. Every model, baselines included, reads it.

The regional models are directed GraphSAGE-style encoders with a pairwise decoder [21]. They are trained only on F1, hiding a fresh random 5% of neurons at every optimization step, and at test time pass messages over the visible part of the target region’s F2 graph. Three non-graph baselines were fitted on F1: a configuration model, logistic regression and gradient-boosted trees (XGBoost) [28]. They score each candidate pair from centroid distance, an F1 estimate of the connection rate between the two cell types, both transmitter labels, innervation similarity, hemisphere match and the granted total. For each region the baselines’ hyperparameters and the strongest baseline family (the “champion”) were chosen on 15 F2 development instances, and the registered comparison used 20 F2 confirmation instances.

### 2.2 Regional models outperform non-graph baselines on a second brain

Table 3 and Fig. 4 give the registered comparison. For each confirmation instance *i*, the model’s advantage is *r_i_* = log_2_(MRR_model_(*i*)*/*MRR_champion_(*i*)), with both MRRs averaged over three training seeds. A region passed if the lower bound of the 95% bias-corrected and accelerated (BCa) bootstrap interval of the mean *r̄* exceeded the region’s noise floor *δ*, and a one-sided sign test over the 20 instances gave *p <* 0.05. *δ* is twice the typical seed-to-seed spread of log_2_ MRR on F1 validation (Methods) and ranged from 0 to 0.03.

**Figure 4:**
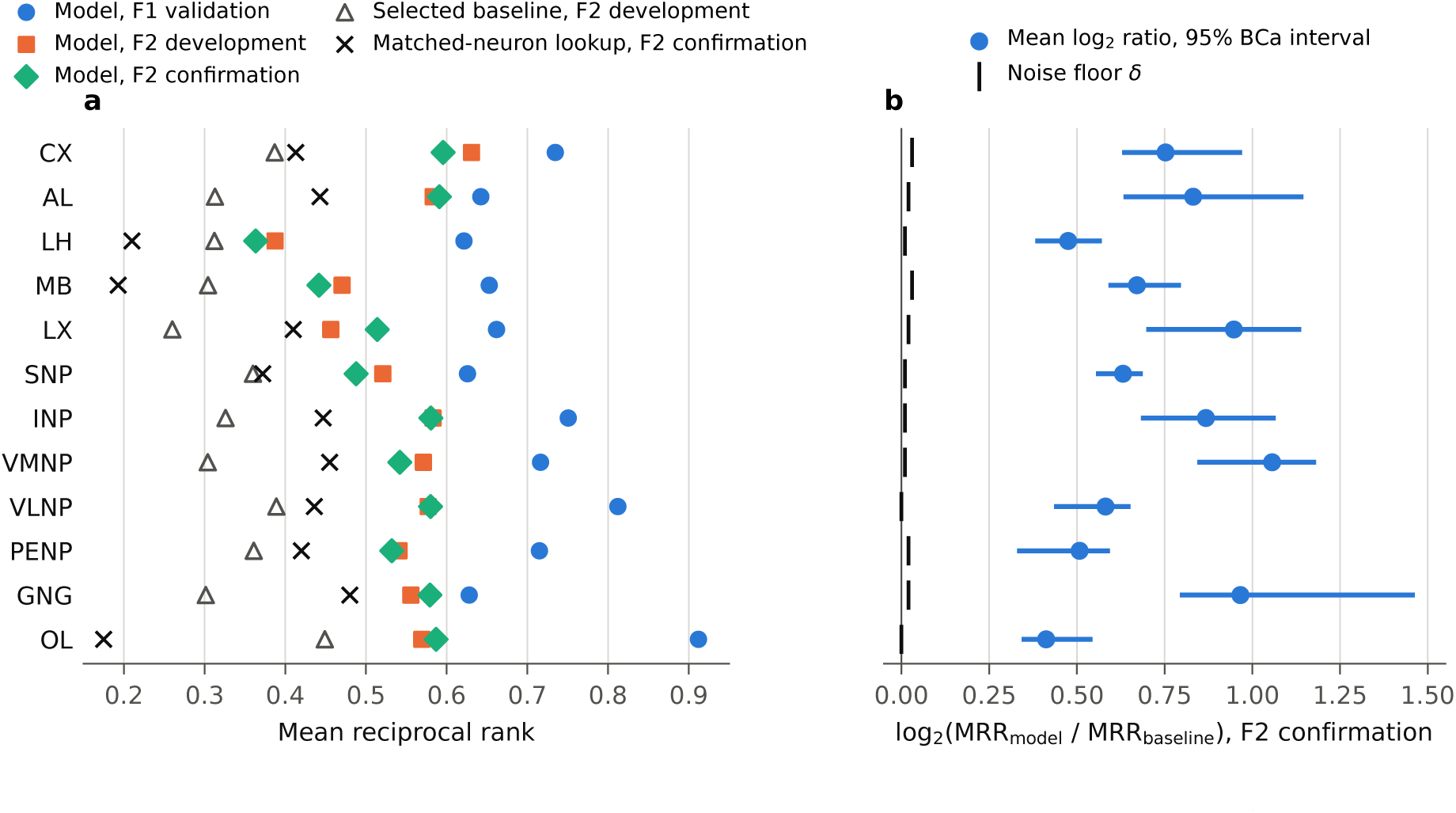
Cross-individual performance per region. **a**, Model MRR on F1 validation (four non-typed instances), F2 development (15 instances) and F2 confirmation (20 instances); the champion baseline’s MRR on F2 development; and, on F2 confirmation, a lookup that copies each masked neuron’s partners from its matched FlyWire neuron (post hoc, see text). **b**, Mean per-instance log_2_ MRR ratio of model to champion on F2 confirmation, with 95% BCa intervals over instances; black ticks mark each region’s noise floor *δ*.

**Table 3:** Registered comparison per region. F1 validation is the model’s MRR over the four non-typed validation instances used for checkpoint selection. F2 columns are three-seed means over 15 development or 20 confirmation instances. The development champion MRR was recomputed from the scored rows of the registered comparison for CX, LH, MB, LX and GNG; for AL (**^ψ^**) it is quoted from the project’s record of that region’s check, and for the others (^‡^) from a separate rescoring in the transfer analysis. *r̄* is the mean per-instance log_2_ MRR ratio against the champion on confirmation, with its 95% BCa interval; wins count instances with *r_i_ >* 0. Confirmation statistics were recomputed from scored rows for CX, LH, MB and LX and match the logged values exactly; for the other regions they are the logged values. Some bounds are printed to four decimals because the log gives no fifth.

| Region | F1 val. | F2 development |  | F2 conf. |  |  |  |  |  |
|---|---|---|---|---|---|---|---|---|---|
|  | Model | Model | Champion | Model | Champion | *δ* | *r̄* [95% BCa] | 2<sup>*r̄*</sup> | Wins |
| CX | 0.735 | 0.631 | 0.387 | 0.596 | XGB | 0.03 | 0.753 [0.629, 0.971] | 1.68 | 19<sup>*</sup> |
| AL | 0.642 | 0.583 | 0.313<sup>†</sup> | 0.591 | XGB | 0.02 | 0.831 [0.633, 1.145] | 1.78 | 18 |
| LH | 0.621 | 0.387 | 0.312 | 0.363 | XGB | 0.01 | 0.475 [0.381, 0.571] | 1.39 | 19 |
| MB | 0.653 | 0.470 | 0.304 | 0.442 | XGB | 0.03 | 0.671 [0.590, 0.7965] | 1.59 | 20 |
| LX | 0.662 | 0.456 | 0.260 | 0.514 | XGB | 0.02 | 0.947 [0.698, 1.139] | 1.93 | 19 |
| SNP | 0.626 | 0.521 | 0.360<sup>‡</sup> | 0.488 | XGB | 0.01 | 0.631 [0.555, 0.688] | 1.55 | 20 |
| INP | 0.751 | 0.583 | 0.326<sup>‡</sup> | 0.580 | XGB | 0.01 | 0.868 [0.683, 1.067] | 1.82 | 20 |
| VMNP | 0.716 | 0.571 | 0.304<sup>‡</sup> | 0.542 | XGB | 0.01 | 1.056 [0.843, 1.182] | 2.08 | 19 |
| VLNP | 0.812 | 0.577 | 0.389<sup>‡</sup> | 0.580 | XGB | 0.00 | 0.5815 [0.4350, 0.6525] | 1.50 | 19 |
| PENP | 0.715 | 0.541 | 0.361<sup>‡</sup> | 0.532 | XGB | 0.02 | 0.508 [0.330, 0.594] | 1.42 | 18 |
| GNG | 0.628 | 0.556 | 0.302 | 0.579 | LR | 0.02 | 0.966 [0.7935, 1.463] | 1.95 | 20 |
| OL | 0.912 | 0.569 | 0.449<sup>‡</sup> | 0.587 | XGB | 0.00 | 0.412 [0.343, 0.544] | 1.33 | 20 |

<sup>\*</sup>Plus one exact tie on an instance with a single scorable edge; sign test *p* = 1.9 × 10<sup>−6</sup> with the tie dropped.

All twelve regions passed. The mean *r̄* ranged from 0.41 in the optic lobe to 1.06 in the ventromedial neuropils, a geometric-mean MRR ratio of 1.33 to 2.08, and every interval lay above *δ* (Fig. 4b). The model won 18 to 20 of 20 instances in every region. In the central complex one instance tied exactly: a typed instance with a single scorable edge, on which the model and both learned baselines ranked the partner first. XGBoost was the champion in eleven regions and logistic regression in the gnathal ganglia. Because decoys are drawn in proportion to visible degree, the configuration model scored near the random expectation (F2 confirmation MRR 0.03–0.04 where these rows are available) except in the mushroom body (0.10). The rule was fixed, with all suites, decoys and *δ* procedures, before any confirmation instance was scored, in an analysis plan the project kept in its own repository; it was not reviewed by a journal, and its deviations are listed in Table 4. The verdicts are reported per region and are not pooled. They are not independent of one another, because about a quarter of graph neurons belong to two or more regions (24.7% in F1, 25.1% in F2). Absolute performance on F2 was lower than on F1 validation (Fig. 4a). Model MRR on the four non-typed F1 validation instances ranged from 0.62 to 0.91; on F2 confirmation it ranged from 0.36

**Table 4:** Status of registered and unregistered analyses.

| Analysis | Status | Notes |
|---|---|---|
| Per-region decision rule on F2 confirmation | Registered; run | All 12 pass. Rule, suites, decoys and *δ* procedure fixed beforehand. Amendments made before any confirmation read are logged: champion chosen on F2 development; decision restricted to single-masked edges; baselines keep 10 of 100 decoys in training; typed F2 masks limited to trained types. |
| Inputs of the tested models | Partly disclosed | Masked innervation computed from complete reconstructions was disclosed on 7 Nov 2026 and the verdicts kept. Position features computed the same way were not disclosed or ablated. |
| Configuration choice | Amended | Screens run before the F2 development suites existed were exempted from the five-configuration rule (amendment of 3 Nov 2026, merged without a recorded review). Each region logged one passing check. |
| *δ* approval order | Disclosed deviation | SNP and VMNP *δ* approved after their confirmation reads; two calibration changes were committed before approval. |
| Order of regional reads | Not recorded in plan | LH and LX were scored on F2 before the central complex verdict was logged, against the project’s own run order. |
| Cross-region transfer matrix | Registered exploratory; run | F2 development only; written up after the confirmatory reads; planned F1 validation version not reported. |
| F3 cross-sex readout | Registered exploratory; run | Model only, no baseline. |
| Confirmation MRR by mask regime | Added after reads | Listed as exploratory on 17 Nov 2026. |
| Post hoc checks | Unregistered; this paper | Matched-neuron lookup, type-mate rule, decoy-shortcut check and transmitter-composition check, run on 17–18 Nov 2026 after all confirmatory reads. |
| Other exploratory analyses | Registered exploratory; not run | Within-F1 edge-holdout rung, degree-free setting, feature ladder, visibility strata, hemisphere-holdout suite, both-masked stratum, boundary split by region count, AP, calibration and Spearman metrics, and topology comparison. Hemisphere-holdout suites were built but not scored; both-masked ranks are stored but not reported, apart from a central complex development readout (Supplementary Table S5). |

(lateral horn) to 0.60 (central complex). The drop was largest in the optic lobe (0.91 to 0.59), lateral horn (0.62 to 0.36) and ventrolateral neuropils (0.81 to 0.58), and smallest in the antennal lobe and gnathal ganglia (about 0.05). F1 validation and F2 differ in the animal and in the reconstruction pipeline. F2’s synapse detector differs from F1’s; for example, its Kenyon cells have about two-thirds as many outgoing connections as FlyWire’s [5]. F2 graphs were also thresholded at calibrated values of 1 to 4 synapses rather than 5. The present data cannot separate these sources of shift. Across the twelve regions, larger drops went with smaller margins over the baseline (Spearman *ρ* = *−*0.68; descriptive).

### 2.3 Direct transfer through matched neurons

The baselines above do not use the wiring of either animal at the level of individual neurons. A more direct form of cross-individual prediction uses the neuron matches BANC provides: most F2 neurons in the central brain have a FlyWire neuron identified as the same cell by morphology search and human review [5]. After the registered comparison, we scored a lookup that ranks each candidate by the number of synapses, within the region, between the FlyWire matches of the masked neuron and the candidate, in the same direction (Methods). It uses no learning and no F2 edges. Outside the optic lobe, 79–100% of confirmation scored edges had both endpoints matched; in the optic lobe, where F2 typing is incomplete, 16% did.

The lookup reached confirmation MRR 0.37–0.48 in nine regions, 0.19–0.21 in the mushroom body and lateral horn, and 0.18 in the optic lobe (Fig. 4a; Supplementary Table S8). Where the champion’s confirmation rows are available, it exceeded the champion in the central complex (0.41 against 0.37) and lateral complex (0.41 against 0.29) and fell below it in the lateral horn and mushroom body; in the other regions it exceeded the champion’s development MRR everywhere except the optic lobe. The matched FlyWire pair shared at least one synapse in the region for 49–76% of scored F2 edges in those nine regions, and for 21% in the mushroom body and 31% in the lateral horn. The regional models beat the lookup in every region. The mean log_2_ ratio ranged from 0.23 in the ventromedial neuropils (95% BCa *−*0.02 to 0.34; 18 of 20 instances) to 2.00 in the optic lobe, and the interval excluded zero in the other eleven regions. Combining the lookup with the type-mate rule described below narrowed the gap but did not close it (log_2_ ratio 0.14–1.45). On typed instances, which hide every type-mate, the lookup beat the model in some regions (for example 0.49 against 0.38 in the ventrolateral neuropils), but each regime has only four instances.

### 2.4 What the margin can and cannot be attributed to

The baselines scored pairwise features built from the same annotations as the models’ inputs, so a passing region shows that the graph model made better use of those inputs and of the visible F2 graph. Several of the inputs carry information about the hidden edges, which limits what “predicting wiring” means here.

#### Inputs computed from the complete reconstruction

The models that passed were trained and scored with a masked neuron’s innervation fractions computed from all of its in-region synapses, including those on its hidden edges; its centroid and spread features were computed the same way. In the eleven F2 regions measured, the synapses on a neuron’s region-graph edges make up a median 12–31% of its in-region synapse ends. The baselines’ innervation-similarity feature reads the same values. This was found after the models were trained, and the regional verdicts were kept, as recorded in the analysis plan’s amendment of 7 November 2026.

Two later measurements bound the innervation part, both on F2 development only. Setting masked innervation to uniform at test time lowered MRR by up to 0.44 in log_2_ units (optic lobe), with little effect where a region has one or two neuropil labels. Retraining all twelve regions with uniform masked innervation in training, selection and scoring changed mean per-instance log_2_ MRR by *−*0.063 in the optic lobe, *−*0.030 in the central complex and less than 0.030 in absolute value elsewhere, and the retrained models remained well above the champions’ development MRR in every region (Supplementary Table S7). The retrained models were not scored on the confirmation suites, the baselines were not rescored with uniform innervation, and the position features were never ablated. The granted total equals the summed weight of the masked neuron’s hidden edges by construction; it is shared with all baselines and is constant across a ranking row, so it can help only in combination with partner features.

#### Cell-type labels

A masked neuron keeps its cell type. F1 types were defined partly from connectivity [3]. F2 neurons received F1 types through their FlyWire matches, which for brain neurons were proposed by morphology search and reviewed visually [5]. The label therefore carries

#### Visible type-mates

In all twelve regions the graph models also read “sibling” features: how many of the masked neuron’s visible same-type neurons (its type-mates) connect to each candidate. No baseline reads any F2 edge apart from the configuration model’s degree counts, and no registered baseline copies the partners of visible type-mates. After the confirmatory reads we scored such a rule on the same suites (Methods). It ranks each candidate by the fraction of the masked neuron’s visible type-mates that connect to it in the same direction. On F2 confirmation it reached MRR 0.09–0.31, and the regional model beat it on all 20 instances in every region (mean log_2_ ratio 1.48–2.58; Supplementary Table S9). It also scored below the champion: in the four regions with champion rows, the champion’s mean log_2_ advantage over the rule was 0.73 in the central complex, where the two split the instances evenly, and 1.43–1.76 in the lateral horn, mushroom body and lateral complex; elsewhere its confirmation MRR lies below the champion’s development MRR. A simple copying rule therefore does not account for the margin. This does not show that type-mate information contributes little inside the graph model, whose sibling features were never ablated.

#### Typed masks

Typed instances hide every neuron of a type, so the masked neuron has no visible type-mates. In an exploratory split of the confirmation scores, added to the plan’s exploratory list after the confirmatory reads, mean model MRR over the twelve regions was 0.506 on typed instances and 0.534–0.544 on the other regimes (Fig. 5). Typed instances scored below the random and spatial regimes in the lateral horn and in the superior, inferior, ventromedial, ventrolateral and periesophageal neuropils. They scored above them in the central complex, antennal lobe, lateral complex, gnathal ganglia and optic lobe, and between them in the mushroom body. Because a type is drawn uniformly among types, most typed masks hide one or two neurons and contribute a median of 101 scorable edges per instance, against thousands for the other regimes. In the four regions with champion rows, the model’s mean log_2_ advantage on typed instances was 0.57, against 0.69–0.88 in the other regimes. With four typed instances per region, this split cannot settle how much the models rely on type-mates.

**Figure 5:**
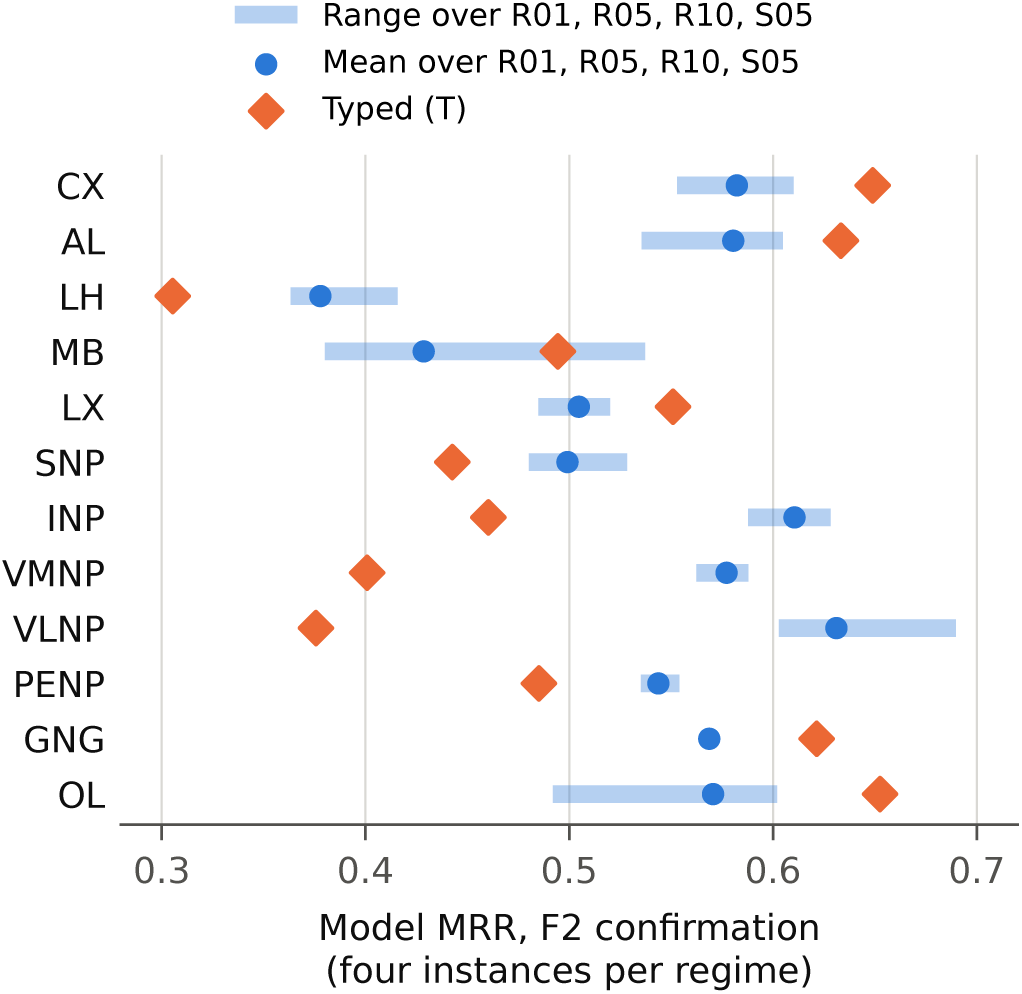
Model MRR on F2 confirmation by mask regime (exploratory split, defined after the confirmatory reads). Orange diamonds: typed masks, which hide every neuron of one cell type. Blue: mean and range over the random (R01, R05, R10) and spatial (S05) regimes. Each regime has four instances per region.

#### Development and confirmation overlap

Development and confirmation suites were disjoint in the cell types that typed masks drew, but random and spatial masks were drawn independently, so 63– 74% of confirmation scored edges had also been hidden in some development instance (Supplementary Table S3). Confirmation therefore re-tests many of the same edges with different neighbours masked. The models’ configurations were chosen on F1 validation screens before any F2 development suite existed, whereas development data were used to tune and choose the baselines, so the overlap can favour only the baselines. It still means the confirmation suite is far from independent of development.

#### A decoy-sampling shortcut

A neuron with zero visible strength on the relevant side can never be drawn as a decoy, so when such a neuron is the true partner, its zero strength marks it as the answer. This affects 0.4–6.2% of confirmation scored edges, most in the mushroom body. On these edges the model scored MRR 0.69–0.83 against 0.34–0.44 for XGBoost, so the cue helps the model more than the baselines. Removing them changed *r̄* by *−*0.017 to +0.013 in the four regions with baseline rows, and no verdict changed (Supplementary Table S10). The effect in the other eight regions (0.5–2.2% of edges) was not computed.

### 2.5 Cross-region transfer does not isolate region-specific wiring rules

To ask whether what a model learns is specific to its region, each regional model was scored on every other region’s F2 development suite (Fig. 6). The plan listed this analysis as exploratory; it was written up after the confirmatory reads, and its row-level output was not deposited, so what follows uses the three-decimal matrices in the project report and has no confidence intervals. On every target region, the region’s own model scored higher than each of the eleven visiting models. Visiting models reached 25–87% of the home model’s MRR (median 53%) and beat the target’s champion baseline in only 30 of 132 region pairs. The home model stayed ahead in every column when its innervation was set to uniform, matching the visitors’ treatment, and when both innervation and position features were removed from home and visitor alike, though then by as little as 0.016 MRR in the lateral horn and mushroom body.

**Figure 6:**
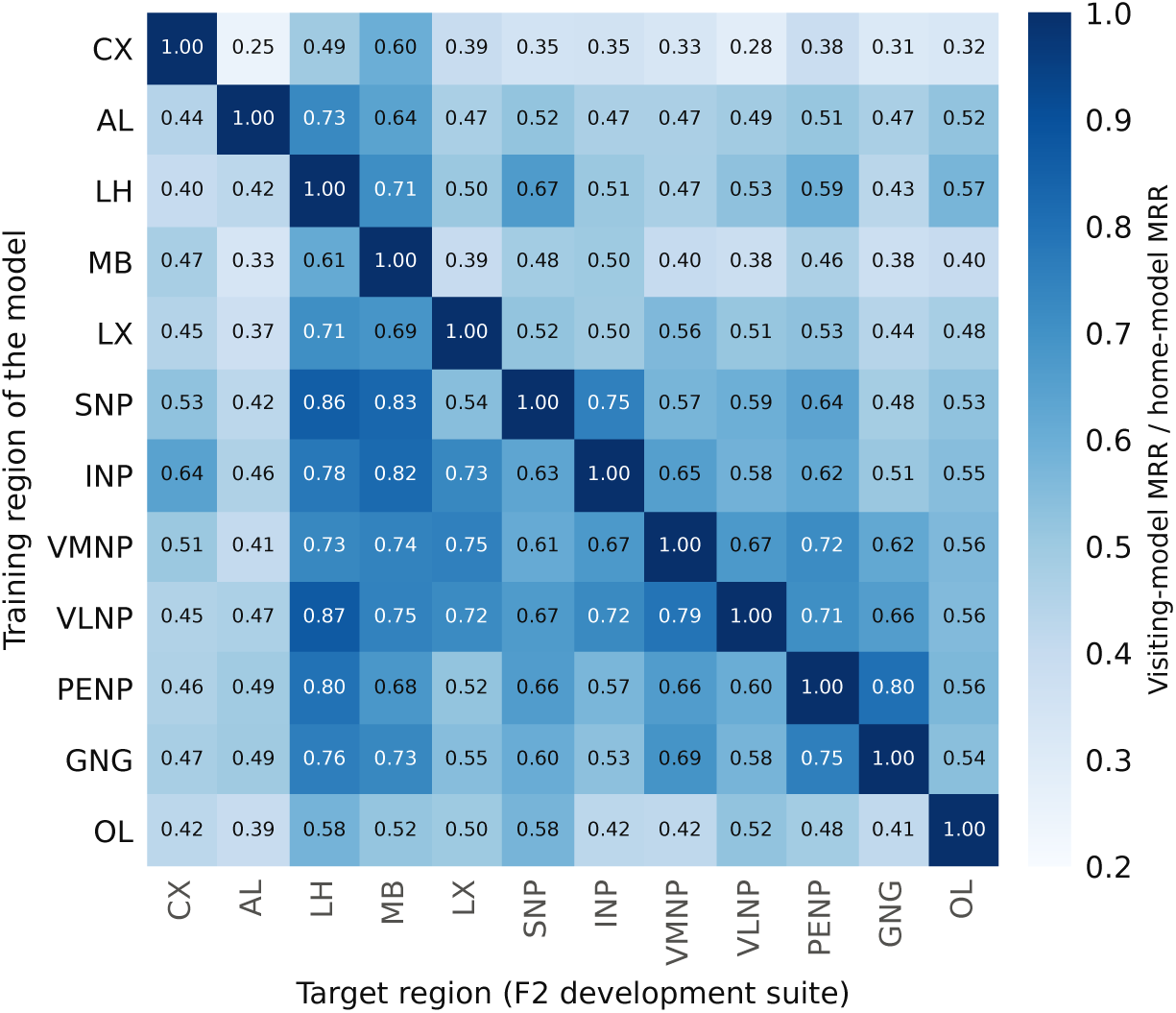
Exploratory cross-region transfer on F2 development suites (computed after the confirmatory reads; no intervals). Each cell is a visiting model’s MRR divided by the target region’s own model’s MRR (three seeds, 15 instances); the diagonal is 1 by definition. Visitors read cell types they did not train on as out of vocabulary and see uniform innervation, so off-diagonal cells mix wiring differences with input mismatch.

Region-specific wiring rules would produce this diagonal dominance, but so would a mismatch of inputs, and the analysis cannot separate the two:

- *Cell-type vocabulary.* A visiting model reads any cell type it did not see in training as “out of vocabulary”, a shared placeholder that also disables its type-mate and type-pair features. Visitors read 18–100% of masked target neurons as out of vocabulary (median 88%); home models read 7–37%.
- *Coordinate frame.* Position features are standardized within each region, so a visitor sees coordinates in a frame it never trained on.
- *Innervation.* Visitors see a uniform innervation profile.

Descriptively, an additive fit with one effect per training region and one per target region (23 parameters for 132 cells) accounts for 82% of the variance in the visiting-to-home ratios. The training-region effect followed the size of each model’s training vocabulary (Spearman *ρ* = 0.89 across twelve models), which is itself confounded with the size of the training graph, and not the model’s home MRR (*ρ* = 0.07). Across region pairs, transfer rose with the overlap of cell-type vocabularies and with the share of neurons the two regions share; these measures correlate at 0.84–0.98 and cannot be separated, and both associations held after removing training-region and target-region effects (permutation *p <* 0.001). Where a visitor knew most of the target’s types, transfer was high: the superior-neuropil model reached 0.83 and 0.86 of home on the mushroom body and lateral horn, and 0.90–0.95 once position and innervation features were removed. These are single cells without intervals.

#### Neurotransmitter composition

Transmitter identity enters the models only as a predicted label per neuron. The regional study did not control for transmitter composition, and no registered analysis tested it. Composition can be computed from the predicted labels, so we checked it post hoc. Similarity of composition between two regions was weakly associated with transfer after removing region effects, and no association remained once vocabulary coverage was also controlled (Spearman *ρ* = 0.06–0.18, permutation *p ≥* 0.10). This null result is not evidence of absence. With twelve regions the test has little power, the labels are predictions [30], transmitter is itself a model input, and the labels disagree between datasets: the F2 label matched the label of the neuron’s FlyWire match for 70.3% of 62,324 matched neurons, and F1 labels most Kenyon cells dopaminergic while F2 labels them cholinergic. The data neither show nor rule out that regional wiring carries information beyond transmitter identity.

The analysis also does not test regularity within a region, reads only F2 development data, and omits the planned within-individual (F1 validation) version.

### 2.6 Exploratory readouts on a male brain

The same models were also scored on 15 masks per region in the male CNS (F3), with no baseline and no selection or tuning on F3. F3 graphs were thresholded at calibrated values of 7–11 synapses.

Pooled over regions and over random and spatial masks, masked neurons of types annotated as isomorphic between the sexes scored a mean per-type MRR of 0.724 when their type was in the model’s training vocabulary and 0.502 when it was not. Male-specific types, which are never in the training vocabulary, scored 0.474, and sexually dimorphic types in the vocabulary scored 0.637. Training status and sex specificity cannot be separated for male-specific types, and without a baseline these numbers describe the models, not a margin (Supplementary Section S8).

## 3 Discussion

The first question has a conditional answer. Given a masked neuron’s cell type, innervation, synapse-position features and total synapse count, and the visible remainder of the second brain’s region graph, regional models trained on one female fly ranked the neuron’s true partners well above chance and above non-graph baselines in all twelve regions, under a rule fixed before the confirmation data were scored. A confirmation MRR of 0.36–0.60 against 100 degree-weighted decoys means the true partner typically ranked among the top few candidates. The result held in regions of very different size and composition, from the 3,219-neuron central complex to the 93,270-neuron optic lobe.

The matched-neuron lookup puts this in context. Copying partners from each neuron’s matched FlyWire counterpart already ranks true partners well in most central-brain regions, and it gives a neuron-level measure of how stereotyped wiring is between these two animals: in nine regions, the matched FlyWire pair shared a synapse for about half to three quarters of scored F2 edges. The regional models did better than the lookup everywhere. Their margin was smallest in the ventromedial neuropils and gnathal ganglia and largest in the optic lobe and mushroom body, where the lookup is weakest. The models do not use the matches directly; they use cell types (assigned in F2 through the matches), position, innervation and the visible F2 graph. Part of their advantage may therefore come from combining type-level regularities with the second animal’s own surrounding wiring, which the lookup ignores.

These results are conditional on what the masked neuron keeps. Its innervation and position features came from its complete reconstruction, its type label came partly from connectivity and, in F2, from matching to F1, and the models also saw the visible wiring of its type-mates. A rule that only copies the type-mates’ partners did poorly, but how much the models’ own type-mate features contribute was not measured. The task is therefore closer to “given who this neuron is and the rest of the circuit, which candidates are its partners” than to predicting wiring from first principles. These caveats limit what kind of knowledge the models hold; they do not change the outcome of the registered comparison.

The second question remains open. A region’s own model beats models trained elsewhere in every region, but visiting models lose most of their type-dependent inputs in a foreign region, their success tracks vocabulary overlap and shared neurons, and in the few cases where a visitor knew the target’s types it came close to the home model. The data cannot distinguish regions that differ mainly in cell-type composition from regions with distinct connection rules that the input mismatch hides. Transmitter composition was not controlled, and a post hoc check had too little power to inform the question. FlyWire neuropils differ in network statistics such as density, reciprocity and motif content [31], and hierarchical modules have been described in the hemibrain [32]; our results neither confirm nor refute that such differences reflect distinct connection rules.

Several designs could separate these explanations. Models trained on a shared, type-agnostic input space, such as morphology and position in a common brain frame, would let visitors and home models read the same inputs. The planned within-individual transfer test would remove dataset shift, and transfer could be measured on subsets of neurons whose types every model has seen. On the first question, controls would bound how much the result depends on identity and degree: an ablation of the models’ own type-mate features, a degree-free setting in which the granted total is withheld, position and innervation features computed only from visible synapses, and the same architecture trained on F2 visible edges alone or left untrained. Several of these were registered as exploratory and never run (Table 4). Whether region-specific wiring rules contribute to specialized functions remains a hypothesis; testing it would need, at minimum, identified wiring patterns that differ between regions and an experimental or modelling link from those patterns to function [33, 34].

### Limitations

- *One pair of animals.* The test uses one training animal and one test animal. The 20 confirmation instances per region mask the same F2 graph and can overlap, so the bootstrap treats dependent units as exchangeable and the intervals are optimistic; the sign test relies on independence less. Instances count equally regardless of size, and typed instances are small.
- *Shared lineage.* F1 and F2 are different animals but not independent annotations: F2’s cell types come from F1 through neuron matches, and its synapse detector and transmitter classifier build on FlyWire-era training data and methods [5].
- *Incomplete data.* Missing F2 edges cannot be told apart from true non-edges. F2 has no lamina, so the optic lobe’s lamina-dependent types were never masked, and only 59.6% of F2 optic-lobe graph neurons have an F1 type.
- *Mushroom body.* Thousands of near-identical Kenyon cells make ranking weak there for every model.
- *Uneven training.* The models trained on fresh simulated gaps with all 100 decoys at every step; the baselines fit once on three fixed gap views with 10 decoys per positive.
- *Sampled decoys.* Evaluation ranks against 100 sampled decoys, not every neuron in the region, so the MRR values do not describe a full partner search.
- *Process.* The departures from the analysis plan are listed in Table 4. Among them: two regions’ noise floors were approved only after their confirmation reads; region configurations were chosen from many F1 validation screens before the F2 suites existed; and the models that passed read hidden-synapse information through innervation and position features.
- *Records.* Confirmation statistics could be recomputed from scored rows for four regions; the other eight rest on the logged values.

## 4 Methods

### 4.1 Datasets

F1 is the FlyWire FAFB v783 release [1] (Zenodo record 10676866): the proofread connection table, with synapses predicted by Buhmann et al. [29], per-neuron neuropil synapse counts and the proofread root identifiers, with annotations from the FlyWire annotation release v3.1.0 [3] (cell type, superclass, class, side, flow and a predicted transmitter per neuron [30]). The table has 139,255 proofread neurons and 54,492,922 synapses and yields 2,700,513 directed pairs with at least five synapses summed across neuropils, matching the published count. The annotation file has 8,840 distinct cell-type strings against a published count of 8,453 types [3]; we did not resolve the difference. F2 is BANC v888 [5]: neuron metadata, the neuropil-resolved edge list and the v2 synapse tables, pinned by checksum. The F2 population is the 155,848 neurons marked proofread or roughly proofread, excluding glia, tracheal and non-neuronal segments. F3 is the male CNS v1.0 [6], restricted to 164,666 neurons of traced or anchor status with a superclass. F1 is a 7-day-old female [2] and F2 a 5–6-day-old female [5], both from crosses of w1118 and Canton-S lines.

### 4.2 Region graphs and harmonization

The twelve regions group FlyWire’s neuropil labels following the standard nomenclature [27]: optic lobe (LA, ME, AME, LO, LOP); lateral complex (BU, LAL, GA); superior neuropils (SLP, SIP, SMP); inferior neuropils (CRE, SCL, ICL, IB, ATL); ventromedial neuropils (VES, EPA, GOR, SPS, IPS); ventrolateral neuropils (AOTU, AVLP, PVLP, PLP, WED); central complex (FB, EB, PB, NO); mushroom body (calyx, pedunculus, vertical and medial lobes); antennal lobe; lateral horn; periesophageal neuropils (SAD, AMMC, FLA, CAN, PRW); and gnathal ganglia. Paired neuropils include both hemispheres. For a region and dataset, synapses on the region’s labels were summed per directed neuron pair; pairs reaching the threshold became edges weighted by that sum, and nodes are the edges’ endpoints. Graphs are directed and never symmetrized. A neuron can belong to several regional graphs, and each regional model sees only its own region’s edges.

F2 and F3 use different synapse detectors, so their thresholds were calibrated per region. For each candidate F2 threshold from 1 to 15, the calibration compared cell-type-pair connection densities between F1 (at threshold 5) and the candidate graph, over type pairs with at least three neurons per side in both datasets, and chose the threshold minimizing the absolute median log_2_ density ratio, breaking ties by rank correlation. The F2 thresholds were frozen before any F2 suite was built. The calibration read complete F2 graphs, including edges later hidden by masks, but no model output. F2 cell types were mapped into F1’s vocabulary in two steps: a neuron took the F1 type of its reviewed FlyWire match, failing that its own BANC type name if that name is an F1 type, and otherwise had no type. About 53% of BANC neurons are matched, with an estimated error rate near 7% [5]. F2 transmitter predictions use eight classes; histamine and tyramine were mapped to “other”. F2 coordinates were reoriented to F1’s axes, and position features were standardized within each dataset’s region graph.

### 4.3 Node features

Each neuron has one-hot superclass and class labels (with an out-of-vocabulary flag), side and flow, a seven-column one-hot transmitter label (six classes and “other”), the standardized centroid and spatial spread of its in-region synapses, and the fraction of its in-region synapses in each of the region’s neuropil labels. Its cell type enters the graph models as a learned 32-dimensional embedding; types with no edge in the region’s F1 training graph, and neurons without a type, share an out-of-vocabulary embedding. In the models that were tested, centroid, spread and innervation were computed from all of a neuron’s in-region synapses, including those of masked neurons. Uniform masked innervation (each label weighted equally) was introduced later and used only in the retrained models.

### 4.4 Masks, suites and decoys

Mask instances cycle through five regimes. R01, R05 and R10 mask a uniformly random 1, 5 or 10% of a region’s neurons; T masks every neuron of one cell type, drawn uniformly among the graph’s typed cell types, in both hemispheres; S05 masks the neurons inside a ball, centred on a random neuron’s centroid, sized to hold 5% of neurons. Each region has 5 F1 validation, 15 F2 development, 20 F2 confirmation and 15 F3 instances, generated once from a fixed seed and pinned by digest. In F2, typed masks draw only types the regional model trains on, and confirmation draws no type that development drew. In the optic lobe, 21 lamina-dependent types (defined on F1 only, from each type’s share of lamina synapses) are never masked but stay in the graph. A mask hides every edge touching a masked neuron, and a hidden edge is scored if exactly one endpoint is masked; edges between two masked neurons are excluded from the decision rule, because counting masked endpoints alone would identify them. For each scored edge, 100 decoys keep the masked endpoint and replace the visible partner with a non-partner of the same region graph, drawn with probability proportional to its visible in-strength (replaced target) or out-strength (replaced source), excluding duplicates and true edges. Decoys were generated once and committed, so every model is scored against identical candidates. The granted total (the masked neuron’s complete in-plus out-strength in the region graph) is given to every model.

### 4.5 Regional models

Each regional model is a two-layer directed GraphSAGE-style encoder with hidden width 128. Each layer combines a neuron’s own representation with separately aggregated incoming and outgoing messages, weighted by log(1 +visible input fraction), with ReLU and dropout of 0.1; ten regions (all but the central complex and optic lobe) add layer normalization, residual connections and branch normalization. A pair decoder, a two-layer perceptron on the concatenated endpoint embeddings and pairwise features, outputs an edge score and a predicted log(1 + synapse count); the same ten regions add a rank-64 bilinear term. Pairwise features are the granted total and the sibling features (the number and fraction of the masked neuron’s visible type-mates connected to the candidate) in all regions, plus, in the same ten regions, cell-type-pair connection densities and pairwise geometry (coordinate differences, distance, hemisphere match and innervation cosine).

Training used F1 only. The training graph is the region’s F1 graph minus every edge touching a neuron masked by any F1 validation instance. At each optimization step (eight per epoch), a fresh random 5% of training neurons was masked and each hidden edge was ranked against 100 decoys from the same sampler. The loss is a softmax cross-entropy over each positive and its decoys plus 0.3 times a smooth-L1 loss on the log synapse count of positives. Cell types were dropped to out of vocabulary with probability 0.1 per step. Optimization used AdamW (weight decay 0.01) with a cosine schedule: learning rate 10^−3^ for 600 epochs in the central complex and optic lobe, 3 *×* 10^−3^ for 600 epochs in the antennal lobe, and 3 *×* 10^−3^ for 200 epochs elsewhere. Every five epochs, the checkpoint was kept if it improved mean MRR on the four non-typed F1 validation instances. Training was deterministic in single precision. Each configuration was trained with seeds 0–4: seeds 0–2 were compared with the baselines, and all five calibrated *δ*. The 60 runs took about 12 GPU-hours on one consumer GPU (estimated from run manifests).

The project team chose each region’s configuration from F1 validation screens run before that region’s F2 development suite existed. A registered check then required the configuration’s three-seed F1 validation MRR to exceed that of the F2-development-selected champion on the same instances; every region passed on its first logged configuration.

### 4.6 Baselines

Logistic regression and XGBoost [28] score a candidate pair from centroid distance, a cross-fitted target encoding of the F1 connection rate of the ordered cell-type pair, both endpoints’ transmitter labels, innervation cosine, hemisphere match and the granted total. The configuration model scores the out-strength of the source times the in-strength of the target, using the granted total for the masked endpoint. The learned baselines were fitted on three fixed F1 gap views (5% of neurons each, from the pinned seed), keeping 10 of the 100 decoys per positive. Hyperparameters were chosen per family on F2 development: 7 regularization strengths for logistic regression and a 24-point grid over depth, number of trees, learning rate and subsampling for XGBoost. The champion was the family with the highest mean F2 development MRR. No baseline reads the edges of the F2 graph apart from the configuration model’s visible strengths.

### 4.7 Metrics and decision rule

For one model, seed and instance, MRR is the mean over scored edges of the reciprocal rank of the true partner among itself and its 100 decoys. Ties are broken pessimistically, so a constant scorer receives 1*/*101. Instance MRRs were averaged over three seeds for both the model and the champion, and *r_i_* = log_2_(MRR_model_(*i*)*/*MRR_champion_(*i*)). A region passed if (i) the lower bound of the 95% BCa bootstrap interval [35] of the mean *r_i_* (10,000 resamples over instances, fixed seed) exceeded *δ*, and (ii) a one-sided exact sign test of *r_i_ >* 0 over the 20 instances (exact zeros dropped) gave *p <* 0.05. Instances are weighted equally regardless of their number of scored edges. *δ* was measured per region before any confirmation read: twice the median, over the five F1 validation instances, of the standard deviation across five seeds of log_2_ MRR, rounded to two decimals, recorded in a tagged file and approved by a second team member; for SNP and VMNP the approval came only after their confirmation reads (Table 4). Confirmation suites were read at milestones declared in advance, and every read was logged. We recomputed the statistics for CX, LH, LX and MB from scored rows with the project’s own code and matched the logged values exactly; for the other eight regions the scored baseline rows were not available for recomputation, and the logged values are reported.

### 4.8 Pre-registration and deviations

The analysis plan was written before data access (18 August 2026) and rewritten for the cross-individual design before any data were scored (29 August 2026). It was kept with a dated amendment log in the project’s version-controlled repository and was not reviewed externally. Its closing revision added the regime split and two limitations after the confirmatory reads. Table 4 lists what was registered, what changed, and what was added or not done.

### 4.9 Post hoc analyses

None of the analyses in this subsection changes a verdict. The regime split, matched-neuron lookup, type-mate rule, decoy-shortcut check and transmitter-composition check were defined and run after every confirmatory read; the transfer matrix was registered as exploratory, and its report was written after the confirmatory reads. *Regime split.* Confirmation MRR by regime is the mean over each regime’s four instances of the three-seed instance MRR, from the committed per-instance scores. *Matched-neuron lookup.* For a scored edge with masked neuron *u* and candidate *v*, the lookup’s score is the number of F1 synapses, summed over the region’s neuropils, from the FlyWire match of *u* to that of *v* (for edges leaving *u*) or the reverse (for edges entering *u*); it is zero if either neuron is unmatched. A binary variant (whether the matched pair is an F1 region-graph edge) scored lower. *Type-mate rule.* The type-mates of *u* are the unmasked neurons of its cell type, as the models read it, with at least one visible edge. The rule scores *v* by the fraction of type-mates with a visible edge to *v* (edges leaving *u*) or from *v* (edges entering *u*); a synapse-weighted variant gave log_2_ ratios within 0.15 of these. Both rules break ties by *v*’s visible strength on the replaced side scaled by 10^−6^, and with no information they reduce to the configuration model. Both were scored with the project’s scoring code on the committed suites and decoys, whose digests were checked, in all twelve regions on F2 development and confirmation, and compared with the models per instance using the same BCa bootstrap. *Decoy shortcut.* We flagged scored edges whose true partner has zero visible strength on the replaced side and, for the four regions whose baseline rows are available, recomputed MRR and the decision statistic without them. *Transfer matrix.* A visiting model was applied to a target region’s F2 development suite through the target’s graph and features, with cell types matched by name, types outside the visitor’s training vocabulary set to out of vocabulary, and innervation set to uniform; each cell is the mean over three seeds and 15 instances. The 132 off-diagonal ratios were decomposed with an additive two-way model, and associations with pair-level properties were tested by Spearman correlation and by permuting region labels (2,000 permutations), before and after removing row and column effects. Regional transmitter composition is the distribution of predicted per-neuron labels over each region’s graph neurons, compared between regions by one minus the Jensen–Shannon distance.

## Data and code availability

FlyWire, BANC and male CNS data are available from their public releases [1, 5, 6]. Code, model configurations, the frozen mask suites with their decoys and digests, noise-floor (*δ*) records, run manifests, the registered verdicts, and the summary tables and scripts behind the figures and post hoc analyses are available at https://github.com/tylerho5/FlyArch under the MIT licence. The analysis plan with its amendment log, the row-level model and baseline scores, the row-level transfer-matrix output and the F3 scores are not in that repository and are available from the authors on reasonable request. The F2 results used BANC v888 files pinned by checksum on 11 October 2026. BANC has since replaced the metadata file in place, filling in the soma side, a model input, for 14,901 neurons; no neuron changed side and cell types, FlyWire matches and the proofread set are unchanged. Scores recomputed from the current file can therefore differ slightly from those reported (antennal-lobe F2 development MRR 0.5818 against 0.5833). The pinned checksums are in the code repository, and the original file is available from the authors on request.

## Author contributions

A.H., T.H. and A.A. contributed equally to this work.

## Supporting information

Supplemental paper

## Acknowledgements

We thank the FlyWire, BANC and FlyEM teams, and the many proofreaders and annotators behind them, for generating and releasing the connectome datasets this work builds on. The authors received no specific funding for this work. AI coding and writing assistants (Claude, Codex) were used during analysis and drafting; the authors reviewed all outputs and take full responsibility for the content.

