## Supplemental paper for "Predicting the synaptic partners of masked neurons across two female *Drosophila* connectomes with per-region graph models"

November 2026

#### Contents

**S1 Data and graph construction** 1

**S2 The central complex in F1** 3

**S3 Mask suites** 6

**S4 Additional regional results** 6

**S5 Masked-innervation diagnosis on F2 development** 7

**S6 Post hoc baselines and decoy shortcut** 8

**S7 Cross-region transfer** 10

**S8 Exploratory readout on the male CNS (F3)** 12

**S9 Attempts to merge the regional models and to route between them** 12
S9.1 Exploratory fusion and routing on F1 . . . 13
S9.2 Pre-registered whole-brain attempt . . . 14

**S10 Pre-registration record** 16

#### S1 Data and graph construction

**Table S1:** Frozen edge thresholds (synapses per directed pair, summed over the region's neuropils). F1 uses 5 in every region. F2 and F3 thresholds were calibrated per region by matching cell-type-pair connection densities to F1 and frozen before any suite on that dataset was built.

| | CX | AL | LH | MB | LX | SNP | INP | VMNP | VLNP | PENP | GNG | OL |
|---|---|---|---|---|---|---|---|---|---|---|---|---|
| F2 | 3 | 4 | 1 | 3 | 2 | 2 | 3 | 2 | 2 | 2 | 3 | 4 |
| F3 | 9 | 11 | 7 | 11 | 8 | 8 | 7 | 8 | 9 | 9 | 11 | 7 |

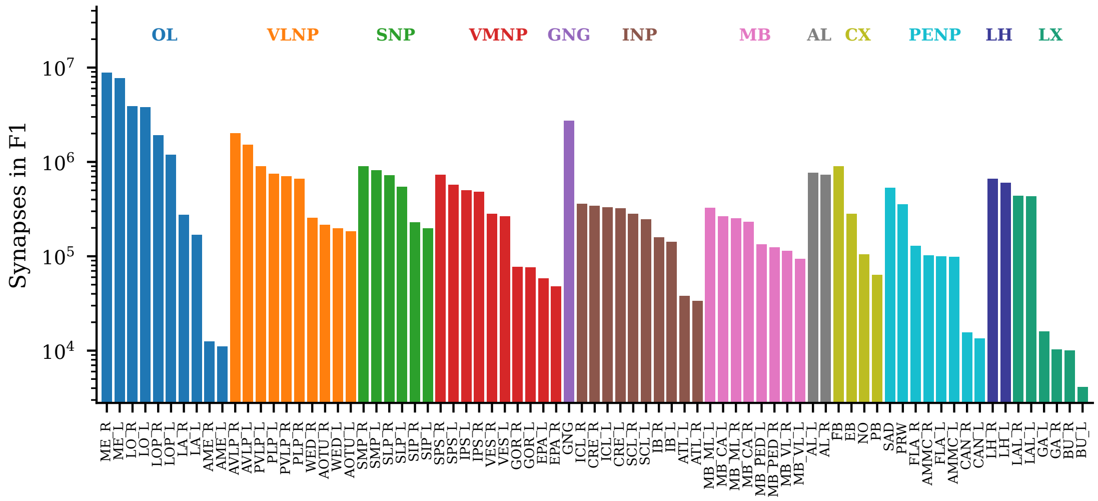


**Figure S1:** F1 synapses in each neuropil label (log scale), grouped and coloured by the region that contains it. Labels outside the twelve regions (OCG and unassigned synapses) are not used.

**Shared neurons.** A neuron belongs to every region graph in which it has an edge. In F1, 33,047 of the 133,844 neurons in any region graph (24.7%) are in two or more; in F2, 32,506 of 129,460 (25.1%). The share of a region's F1 graph neurons that also appear in another region's graph is 99.7% (LX), 98.9% (INP), 96.8% (VMNP), 93.8% (LH), 90.3% (VLNP), 88.2% (PENP), 84.5% (SNP), 73.3% (MB), 65.1% (GNG), 54.1% (CX), 31.1% (AL) and 8.5% (OL). Each regional model reads only its own region's graph, so shared neurons are not an input route, but they make the twelve regional verdicts statistically dependent.

**F2 cell types and dependence on F1.** F2 neurons take the F1 cell type of their reviewed FlyWire match, else their own BANC type name if it is an F1 type. BANC's matches were proposed by NBLAST and reviewed visually, with connectivity used for some classes; about 53% of BANC neurons are matched, with an estimated error rate near 7% [1]. F2's superclass and class labels were mapped to the F1 value most common among matched neurons. BANC's synapse detector and neurotransmitter classifier were trained with FlyWire-era data and methods [1, 2]. F1 and F2 are therefore distinct animals but not independent annotations.

**Neurotransmitter labels.** Both datasets supply one predicted transmitter label per neuron. F1 uses six classes; F2 uses eight, and histamine and tyramine were mapped to "other". For 62,324 F2 neurons with a FlyWire match, the F2 label agrees with the F1 label of the match in 70.3% (54.9% for central-brain neurons, 88.0% for optic-lobe neurons). In F1, 5,172 of 5,177 Kenyon cells carry a predicted dopaminergic label, whereas F2 labels matched Kenyon cells cholinergic. Table S2 gives F1 composition by region.

**Table S2:** Predicted transmitter composition of F1 region graphs (fraction of graph neurons). The mushroom body's dopamine fraction reflects the Kenyon-cell labels noted in the text.

| Region | ACh | Glu | GABA | DA | 5-HT | OA |
|---|---|---|---|---|---|---|
| CX | .587 | .220 | .122 | .024 | .045 | .001 |
| AL | .647 | .063 | .115 | .009 | .165 | .002 |
| LH | .567 | .267 | .153 | .009 | .004 | .001 |
| MB | .205 | .109 | .044 | .636 | .004 | .002 |
| LX | .543 | .202 | .228 | .018 | .006 | .003 |
| SNP | .520 | .294 | .083 | .092 | .010 | .001 |
| INP | .470 | .224 | .113 | .184 | .006 | .002 |
| VMNP | .567 | .208 | .206 | .008 | .006 | .004 |
| VLNP | .691 | .138 | .138 | .027 | .004 | .002 |
| PENP | .580 | .129 | .237 | .008 | .037 | .008 |
| GNG | .621 | .127 | .201 | .013 | .034 | .005 |
| OL | .676 | .174 | .146 | .001 | .003 | .001 |

#### S2 The central complex in F1

The central complex (CX; ellipsoid body EB, fan-shaped body FB, protocerebral bridge PB and noduli NO) is the smallest region and was the first one the project studied. The figures below describe its F1 graph (3,219 neurons, 70,830 edges at five or more synapses) and its validation masks. They are descriptive and are drawn from the F1 release alone.

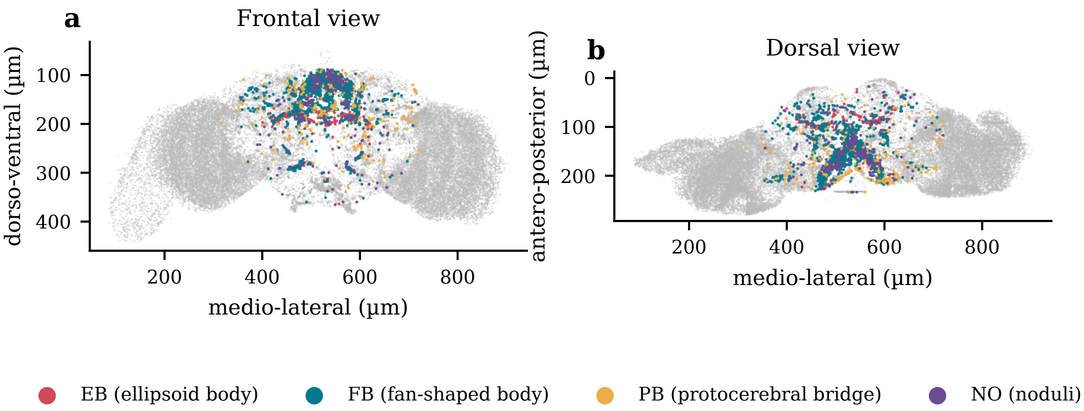


**Figure S2:** Central complex neurons within the F1 brain, coloured by each neuron's primary CX neuropil, over a random sample of 40,000 F1 neurons (grey). **a**, frontal view; **b**, dorsal view.

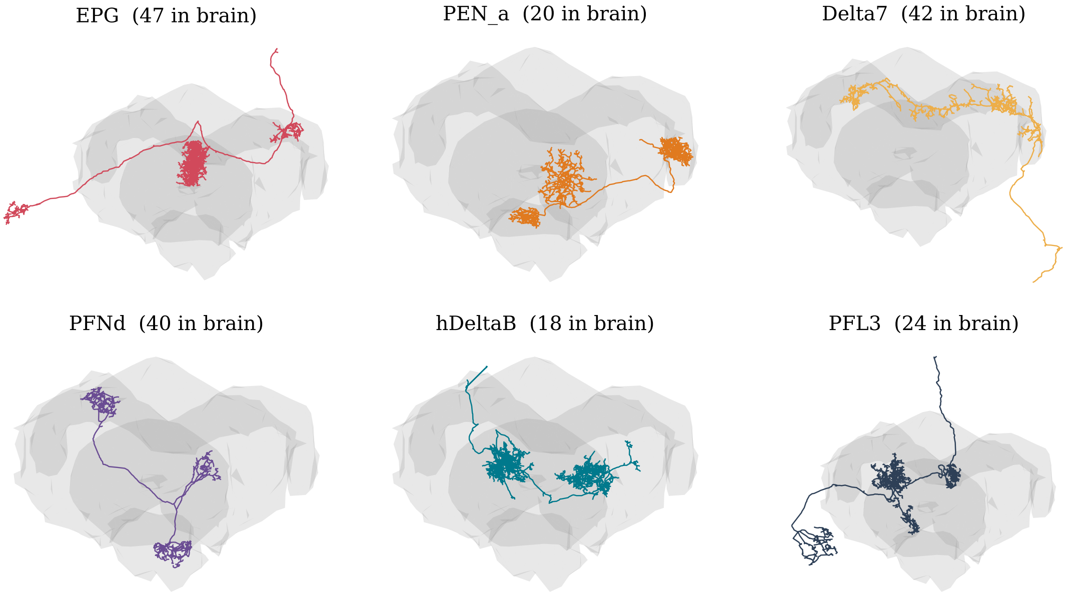


**Figure S3:** Six central complex cell types, one example neuron each, rendered from public FlyWire v783 skeletons inside the brain mesh. Numbers give the type's neuron count in the F1 annotation release (EPG 47, PEN_a 20, Delta7 42, PFNd 40, hDeltaB 18, PFL3 24).

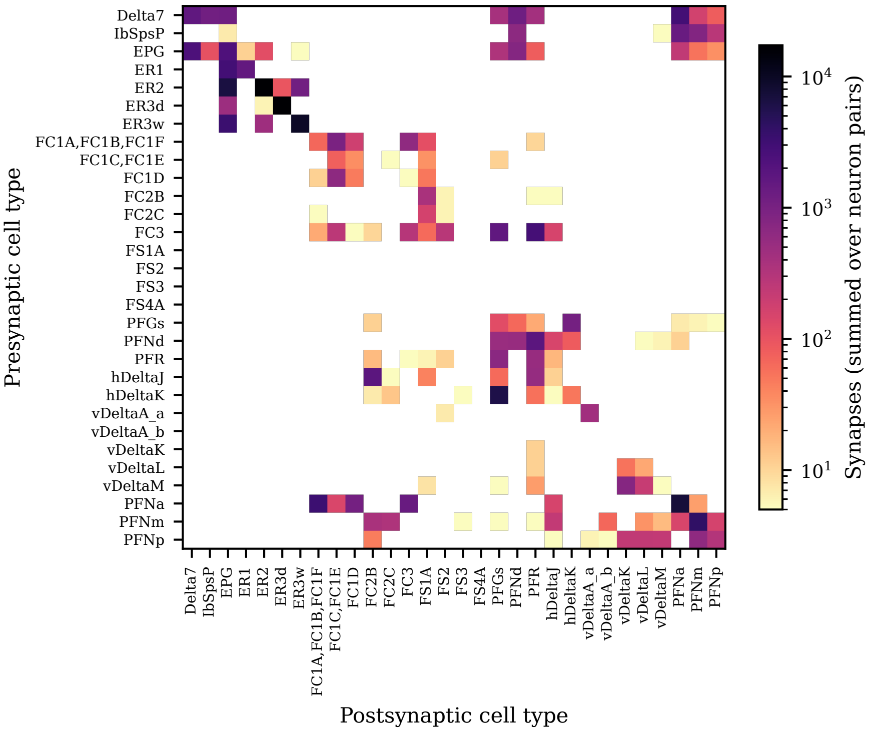


**Figure S4:** Central complex connectivity between the 30 most numerous cell types: synapses summed over all neuron pairs of each presynaptic (rows) and postsynaptic (columns) type, within the CX region graph (log colour scale).

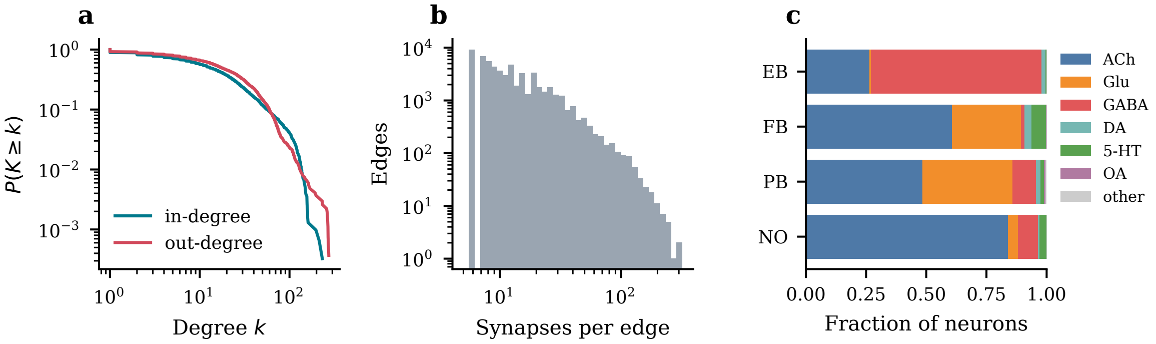


**Figure S5:** Central complex F1 graph statistics. **a**, complementary cumulative distributions of in- and out-degree; **b**, synapses per edge (edges have at least five); **c**, predicted transmitter composition of neurons by primary neuropil.

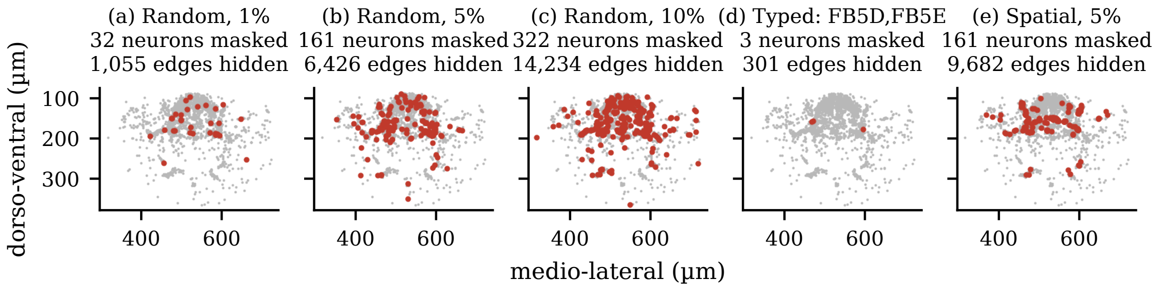


**Figure S6:** The five central complex F1 validation instances, one per regime. Red: masked neurons; grey: other CX neurons (frontal projection). The typed instance masks one cell type of three neurons.

#### S3 Mask suites

Instances cycle R01, R05, R10, T, S05 with seeds offset per split (F1 validation 0–4, F2 development 1000–1014, F2 confirmation 2000–2019, F3 3000–3014). Typed instances draw a cell type uniformly among the graph's typed types, so most hide one or two neurons. On F2 confirmation they contribute between 1 and 2,176 scored edges (median 101), against medians of 2,780, 12,206, 23,634 and 8,614 for R01, R05, R10 and S05.

**Table S3:** Overlap between F2 development and confirmation suites, and decoy-sampling shortcut rows. "Masked in dev" is the share of confirmation-masked neurons masked in some development instance. "Edges hidden in dev" is the share of confirmation scored edges that were hidden edges in some development instance. "Zero-strength partner" is the share of confirmation scored edges whose true partner has zero visible strength on the replaced side and so can never appear as a decoy.

| Region | Scored edges | Masked in dev | Edges hidden in dev | Zero-strength partner |
|---|---|---|---|---|
| CX | 107,913 | 49.5% | 73.2% | 0.42% |
| AL | 111,995 | 47.7% | 63.1% | 0.69% |
| LH | 259,717 | 46.5% | 70.7% | 0.38% |
| MB | 44,375 | 48.1% | 69.0% | 6.24% |
| LX | 101,147 | 48.2% | 74.3% | 0.81% |
| SNP | 302,535 | 48.4% | 72.5% | 0.67% |
| INP | 188,948 | 46.5% | 68.8% | 2.11% |
| VMNP | 380,798 | 51.3% | 71.4% | 0.47% |
| VLNP | 640,565 | 50.3% | 71.7% | 0.46% |
| PENP | 139,668 | 45.3% | 70.0% | 0.92% |
| GNG | 195,471 | 48.1% | 71.0% | 1.02% |
| OL | 913,788 | 45.9% | 67.3% | 2.22% |

#### S4 Additional regional results

**Table S4:** MRR and Recall@10 of the model and all three baselines, for the four regions whose scored baseline rows were available (CX, LH, MB, LX). Three-seed means over 15 development or 20 confirmation instances.

| Region | Suite | Model | XGBoost | Logistic regression | Configuration |
|---|---|---|---|---|---|
| CX | dev | .631 / .907 | .387 / .679 | .367 / .659 | .032 / .052 |
| CX | confirm | .596 / .883 | .368 / .629 | .362 / .623 | .037 / .059 |
| LH | dev | .387 / .727 | .312 / .622 | .310 / .612 | .036 / .058 |
| LH | confirm | .363 / .698 | .264 / .567 | .265 / .570 | .036 / .060 |
| MB | dev | .470 / .700 | .304 / .495 | .264 / .468 | .099 / .172 |
| MB | confirm | .442 / .670 | .290 / .504 | .224 / .455 | .100 / .171 |
| LX | dev | .456 / .749 | .260 / .508 | .252 / .492 | .038 / .065 |
| LX | confirm | .514 / .790 | .293 / .502 | .283 / .477 | .033 / .053 |
| GNG | dev | .556 / .856 | .299 / .588 | .302 / .587 | .038 / .057 |

Each cell: MRR / Recall@10. Random ranking against 100 decoys gives MRR .051 and Recall@10 .099.

**Table S5:** Central complex, F2 development, pooled over scored edges by stratum (three-seed mean per edge). "Both" masked edges are excluded from the decision rule and shown for reference. "Unmatched" means the masked neuron's type was not in the model's training vocabulary.

| | | | XGBoost | | Model | |
|---|---|---|---|---|---|---|
| Axis | Stratum | Edges | MRR | R@10 | MRR | R@10 |
| Masked endpoints | one | 73,336 | .347 | .614 | .585 | .894 |
| | both | 5,105 | .514 | .769 | .982 | .998 |
| Direction | masked source | 37,242 | .344 | .614 | .590 | .901 |
| | masked target | 36,094 | .351 | .614 | .579 | .886 |
| Type | matched | 73,270 | .348 | .614 | .585 | .894 |
| | unmatched | 66 | .253 | .546 | .335 | .490 |

**Table S6:** Model MRR on F2 confirmation by mask regime (four instances each; exploratory split defined after the confirmatory reads). The last two rows give, for the four regions with available baseline rows (CX, LH, MB, LX), the mean champion MRR and the mean per-instance $\log_2$ ratio by regime.

| Region | R01 | R05 | R10 | T | S05 |
|---|---|---|---|---|---|
| CX | .578 | .610 | .588 | .649 | .553 |
| AL | .605 | .595 | .587 | .633 | .536 |
| LH | .368 | .363 | .364 | .305 | .416 |
| MB | .405 | .392 | .380 | .494 | .537 |
| LX | .517 | .497 | .485 | .551 | .520 |
| SNP | .480 | .491 | .497 | .443 | .528 |
| INP | .628 | .614 | .612 | .460 | .588 |
| VMNP | .588 | .581 | .578 | .401 | .562 |
| VLNP | .623 | .609 | .603 | .376 | .690 |
| PENP | .537 | .554 | .548 | .485 | .535 |
| GNG | .570 | .572 | .567 | .621 | .567 |
| OL | .590 | .602 | .598 | .653 | .492 |
| Mean of 12 | .541 | .540 | .534 | .506 | .544 |
| Champion, mean of 4 | .288 | .282 | .280 | .382 | .287 |
| $\log_2$ ratio, mean of 4 | .702 | .713 | .690 | .570 | .883 |

### S5 Masked-innervation diagnosis on F2 development

The regional models that passed the registered comparison read masked neurons' innervation computed from all of their in-region synapses. Table S7 summarizes a later diagnosis on F2 development suites (pre-registered as exploratory; it selects nothing and changes no verdict). "Uniform at test" rescored the same models with masked innervation set to uniform. The retrained cohort was trained, selected and scored with uniform masked innervation (seeds 0–4; three-seed means shown). "Union graph" and "A" come from the whole-brain attempts (Section S9). In GNG the region has a single neuropil label, so uniform and real innervation coincide and the retrained cohort reproduces the original exactly. No confirmation suite was read for the retrained cohort, and no baseline was rescored with uniform innervation.

**Table S7:** F2 development MRR (mean of 15 instances, three seeds) by model and input treatment. $\Delta$ is the mean per-instance $\log_2$(retrained / original). Union-graph scores are missing where the union graph lacked some of the region's F2 neurons when this was computed.

| | Original models | | Retrained, uniform | | | |
|---|---|---|---|---|---|---|
| Region | As scored | Uniform at test | Region graph | Union graph | $\Delta$ | A |
| CX | .631 | .592 | .616 | .492 | −.030 | .556 |
| AL | .583 | .568 | .596 | .520 | +.030 | .477 |
| LH | .387 | .389 | .390 | – | +.003 | .276 |
| MB | .470 | .464 | .482 | .409 | +.024 | .422 |
| LX | .456 | .428 | .453 | .352 | −.010 | .404 |
| SNP | .521 | .437 | .512 | – | −.028 | .423 |
| INP | .583 | .517 | .581 | .527 | −.002 | .522 |
| VMNP | .571 | .539 | .567 | – | −.016 | .483 |
| VLNP | .577 | .534 | .581 | – | +.012 | .493 |
| PENP | .541 | .516 | .548 | – | +.026 | .460 |
| GNG | .556 | .556 | .556 | .497 | .000 | .501 |
| OL | .569 | .421 | .546 | .378 | −.063 | .576 |

#### S6 Post hoc baselines and decoy shortcut

These analyses were defined and run on 17–18 November 2026, after every confirmatory read. They are not in the pre-registration and change no verdict.

**Matched-neuron lookup.** For a scored edge with masked neuron $u$ and candidate $v$, the lookup scores $v$ by the number of F1 synapses, summed over the region's neuropils, from the FlyWire match of $u$ to the FlyWire match of $v$ (for edges leaving $u$), or from $v$'s match to $u$'s (for edges entering $u$). Matches are BANC's reviewed FlyWire matches; several BANC neurons can share one FlyWire match. The score is zero if either neuron is unmatched, and ties are broken by $v$'s visible F2 strength on the replaced side, scaled by $10^{-6}$. A binary variant (whether the matched pair is an edge of the F1 region graph) scored lower in every region. The lookup uses no learning and no F2 edges, but it uses neuron-level identity that the regional models and baselines do not receive.

Restricted to scored edges whose two endpoints are matched, the lookup's MRR changes little outside the optic lobe (0.21–0.49) and reaches 0.57 in the optic lobe. Pooled over such edges in the four regions with baseline rows, the model, the lookup and XGBoost scored 0.578, 0.466 and 0.348 in CX; 0.496, 0.454 and 0.243 in LX; 0.372, 0.225 and 0.263 in LH; and 0.388, 0.207 and 0.264 in MB. By regime, the lookup beat the model on some typed instances (2 of 4 in INP, VMNP, PENP, LH and LX; 3 of 4 in VLNP) and on at most 2 of 16 non-typed instances in any region (LX; none elsewhere).

**Type-mate heuristic.** For a scored edge with masked neuron $u$, the rule's score for candidate $v$ is the fraction of $u$'s type-mates with a visible edge to $v$ (edges leaving $u$) or from $v$ (edges entering $u$). A type-mate is an unmasked neuron of $u$'s cell type with at least one visible edge, using the same crosswalked, training-vocabulary type identifiers the models read. Ties are broken by $v$'s visible strength on the replaced side divided by the maximum in the instance, times $10^{-6}$. With no type or no visible type-mate, only the tie-breaker ranks; the rule then reproduces the configuration model's MRR. A synapse-weighted variant (mean visible synapse count from type-mates) gave $\log_2$ ratios within 0.15 of those shown. Negatives were regenerated with the project's sampler and matched the

**Table S8:** Matched-neuron lookup (L) on F2 (instance means; 15 development, 20 confirmation instances). The $\log_2$ ratio is the mean per-instance $\log_2(\mathrm{MRR}_{\mathrm{model}}/\mathrm{MRR}_{\mathrm{L}})$ on confirmation with a 95% BCa interval; wins out of 20. "Both matched" is the share of confirmation scored edges whose two endpoints have FlyWire matches; "F1 synapse" is the share whose matched FlyWire pair shares at least one synapse in the region. "L + type-mate" ranks by the lookup and breaks its ties by the type-mate rule.

| | L | | | | | L + type-mate | | | |
|---|---|---|---|---|---|---|---|---|---|
| Region | Dev | Conf | Model | $\log_2$(model/L) [95% BCa] | Wins | Conf | $\log_2$ ratio | Both matched | F1 synapse |
| CX | .410 | .413 | .596 | 0.77 [0.34, 1.97] | 20 | .441 | 0.68 | 99.6% | 69.7% |
| AL | .463 | .443 | .591 | 0.45 [0.31, 0.64] | 19 | .461 | 0.39 | 97.6% | 76.4% |
| LH | .224 | .210 | .363 | 0.78 [0.59, 1.02] | 18 | .229 | 0.66 | 96.6% | 31.4% |
| MB | .199 | .193 | .442 | 1.16 [1.04, 1.30] | 20 | .204 | 1.09 | 78.8% | 20.7% |
| LX | .437 | .410 | .514 | 0.68 [0.09, 1.92] | 16 | .428 | 0.62 | 98.4% | 60.9% |
| SNP | .392 | .372 | .488 | 0.40 [0.31, 0.52] | 19 | .393 | 0.33 | 97.9% | 49.5% |
| INP | .484 | .447 | .580 | 0.37 [0.18, 0.46] | 18 | .496 | 0.22 | 96.8% | 54.9% |
| VMNP | .428 | .455 | .542 | 0.23 [−0.02, 0.34] | 18 | .485 | 0.14 | 95.8% | 55.6% |
| VLNP | .385 | .436 | .580 | 0.39 [0.13, 0.53] | 17 | .474 | 0.27 | 89.7% | 52.4% |
| PENP | .380 | .420 | .532 | 0.36 [0.16, 0.47] | 18 | .462 | 0.21 | 96.9% | 48.8% |
| GNG | .493 | .480 | .579 | 0.28 [0.20, 0.39] | 19 | .517 | 0.18 | 95.9% | 58.5% |
| OL | .140 | .175 | .587 | 2.00 [1.58, 2.27] | 19 | .235 | 1.45 | 16.3% | 9.9% |

committed digests for every instance.

Where per-instance champion rows are available, the champion's mean $\log_2$ advantage over the rule on confirmation was 0.73 [0.17, 1.88] in CX (10 of 20 instances), 1.76 [1.47, 2.40] in LH (20 of 20), 1.48 [0.94, 2.52] in LX (20 of 20) and 1.43 [0.91, 2.38] in MB (19 of 20). On the sixteen non-typed instances, the rule's $\log_2$ ratio to the champion was +0.07 in CX (rule ahead on 10 of 16) and between −0.80 and −1.48 in the other three regions. Pooled over scored edges whose masked neuron has a visible type-mate, the rule's MRR in CX was 0.433 against 0.597 for the model and 0.358 for XGBoost. In LH, LX and MB it was 0.11–0.16, below every learned baseline. The rule's scores take few distinct values before tie-breaking, which may lower its MRR, and stronger copying rules (for example, a learned combination of type-mate overlap and degree, or the models' sibling features alone) were not tried.

**Decoy shortcut.** A scored edge is flagged when its true partner has zero visible strength on the replaced side, so that the degree-weighted sampler can never draw it as a decoy (Table S3). Table S10 recomputes the comparison without the flagged edges for the four regions whose scored baseline rows are available.

**Table S9:** Type-mate heuristic (H) on F2. MRR is the mean over instances (15 development, 20 confirmation). Model MRR is the three-seed regional model. The $\log_2$ ratio is the mean per-instance $\log_2(\mathrm{MRR}_{\mathrm{model}}/\mathrm{MRR}_{\mathrm{H}})$ on confirmation, with a 95% BCa interval; the model won all 20 instances in every region. "With type-mate" is the share of confirmation scored edges whose masked neuron has at least one visible type-mate. "T" columns are the four typed confirmation instances, where no type-mate is visible.

| | Development | | Confirmation | | | | |
|---|---|---|---|---|---|---|---|
| Region | H | Model | H | Model | $\log_2$(model/H) [95% BCa] | With type-mate | T: H / model |
| CX | .310 | .631 | .311 | .596 | 1.48 [0.89, 2.62] | 87.1% | .029 / .649 |
| AL | .221 | .583 | .187 | .591 | 2.11 [1.59, 3.10] | 94.0% | .038 / .633 |
| LH | .084 | .387 | .086 | .363 | 2.23 [1.95, 2.84] | 87.3% | .034 / .305 |
| MB | .123 | .470 | .122 | .442 | 2.11 [1.62, 2.97] | 78.9% | .028 / .494 |
| LX | .128 | .456 | .121 | .514 | 2.42 [1.98, 3.25] | 91.0% | .022 / .551 |
| SNP | .137 | .521 | .133 | .488 | 2.11 [1.74, 2.73] | 72.3% | .034 / .443 |
| INP | .224 | .583 | .221 | .580 | 1.76 [1.37, 2.41] | 86.5% | .031 / .460 |
| VMNP | .171 | .571 | .166 | .542 | 2.00 [1.68, 2.67] | 89.7% | .032 / .401 |
| VLNP | .182 | .577 | .176 | .580 | 2.04 [1.68, 2.57] | 87.3% | .033 / .376 |
| PENP | .201 | .541 | .188 | .532 | 1.90 [1.46, 2.64] | 84.1% | .026 / .485 |
| GNG | .222 | .556 | .210 | .579 | 1.78 [1.37, 2.57] | 79.6% | .082 / .621 |
| OL | .130 | .569 | .122 | .587 | 2.58 [2.16, 3.18] | 62.1% | .026 / .653 |

**Table S10:** Sensitivity of the registered comparison to removing flagged edges (F2 confirmation). MRR columns are instance means with all edges and with flagged edges removed. "Flagged only" pools the flagged edges over seeds. $\bar{r}$ is the mean per-instance $\log_2$ ratio of model to XGBoost with its 95% BCa interval; the number of instances won (19, 19, 19 and 20) is the same before and after removal.

| | | MRR, all → removed | | Flagged only | $\bar{r}$ [95% BCa] | |
|---|---|---|---|---|---|---|
| Region | Flagged | Model | XGBoost | Model / XGBoost | All edges | Flagged removed |
| CX | 0.42% | .596 → .592 | .368 → .369 | .825 / .419 | 0.753 [0.629, 0.971] | 0.735 [0.616, 0.896] |
| LH | 0.38% | .363 → .362 | .264 → .264 | .687 / .342 | 0.475 [0.381, 0.571] | 0.468 [0.374, 0.564] |
| LX | 0.81% | .514 → .511 | .293 → .285 | .791 / .439 | 0.947 [0.698, 1.139] | 0.961 [0.715, 1.144] |
| MB | 6.24% | .442 → .423 | .290 → .279 | .691 / .369 | 0.671 [0.590, 0.7965] | 0.654 [0.575, 0.780] |

### S7 Cross-region transfer

Each regional model (seeds 0–2) was scored on every other region's 15 F2 development instances. For a visiting model, the target's features were re-encoded: cell types matched by name (the twelve regional schemas share one F1 vocabulary of 8,840 type names), types outside the visitor's training vocabulary read as out of vocabulary, innervation set to uniform, and position features left in the target's own standardized frame. The visitor passes messages over the target's visible F2 graph and ranks against the target's committed decoys. Diagonal cells were also scored with uniform innervation and with innervation and position features removed. The analysis is listed as exploratory in the pre-registration. Its report was committed after the twelve confirmatory reads, and its row-level output is not in the repository. All numbers here are from the report's three-decimal matrices.

**Decomposition.** An additive model with a training-region and a target-region effect explains 82.3% of the variance of the 132 off-diagonal visiting-to-home ratios (43.3% for training region alone, 38.6% for target region alone; 84.3% for log ratios). The training-region effect follows the size of the

**Table S11:** Visiting-model MRR on F2 development (rows: training region; columns: target region). Diagonal: home model.

| Model | CX | AL | LH | MB | LX | SNP | INP | VMNP | VLNP | PENP | GNG | OL |
|---|---|---|---|---|---|---|---|---|---|---|---|---|
| CX | **.631** | .144 | .192 | .281 | .178 | .182 | .205 | .189 | .163 | .207 | .173 | .183 |
| AL | .279 | **.583** | .281 | .301 | .215 | .271 | .273 | .266 | .282 | .278 | .262 | .298 |
| LH | .252 | .247 | **.387** | .335 | .230 | .347 | .295 | .269 | .305 | .322 | .239 | .327 |
| MB | .298 | .195 | .238 | **.470** | .180 | .252 | .291 | .228 | .220 | .249 | .208 | .226 |
| LX | .281 | .215 | .274 | .323 | **.456** | .271 | .289 | .321 | .296 | .287 | .244 | .272 |
| SNP | .331 | .246 | .332 | .391 | .247 | **.521** | .439 | .326 | .343 | .345 | .268 | .300 |
| INP | .404 | .266 | .301 | .385 | .334 | .329 | **.583** | .372 | .338 | .334 | .283 | .311 |
| VMNP | .319 | .236 | .282 | .349 | .343 | .319 | .392 | **.571** | .389 | .388 | .344 | .320 |
| VLNP | .284 | .271 | .337 | .351 | .329 | .347 | .422 | .450 | **.577** | .383 | .366 | .316 |
| PENP | .289 | .287 | .308 | .320 | .237 | .344 | .332 | .378 | .348 | **.541** | .445 | .320 |
| GNG | .300 | .288 | .294 | .343 | .249 | .315 | .311 | .395 | .333 | .408 | **.556** | .308 |
| OL | .267 | .228 | .225 | .243 | .227 | .301 | .248 | .238 | .301 | .262 | .228 | **.569** |

**Table S12:** Share of masked target neurons (single-masked scored edges) whose cell type the row's model reads as out of vocabulary.

| Model | CX | AL | LH | MB | LX | SNP | INP | VMNP | VLNP | PENP | GNG | OL |
|---|---|---|---|---|---|---|---|---|---|---|---|---|
| CX | **0.12** | 0.99 | 0.99 | 0.96 | 0.86 | 0.94 | 0.90 | 0.90 | 0.98 | 0.97 | 0.98 | 0.93 |
| AL | 0.93 | **0.10** | 0.84 | 0.95 | 0.90 | 0.97 | 0.90 | 0.96 | 0.92 | 0.90 | 0.95 | 1.00 |
| LH | 1.00 | 0.75 | **0.17** | 0.61 | 0.94 | 0.63 | 0.72 | 0.94 | 0.90 | 0.94 | 0.96 | 1.00 |
| MB | 0.83 | 0.85 | 0.69 | **0.20** | 0.57 | 0.72 | 0.61 | 0.86 | 0.89 | 0.94 | 0.97 | 1.00 |
| LX | 0.81 | 0.96 | 0.96 | 0.96 | **0.15** | 0.95 | 0.87 | 0.80 | 0.89 | 0.85 | 0.94 | 1.00 |
| SNP | 0.62 | 0.81 | 0.26 | 0.18 | 0.60 | **0.08** | 0.38 | 0.80 | 0.69 | 0.83 | 0.91 | 0.99 |
| INP | 0.40 | 0.87 | 0.44 | 0.24 | 0.23 | 0.45 | **0.11** | 0.56 | 0.63 | 0.82 | 0.87 | 0.92 |
| VMNP | 0.78 | 0.87 | 0.88 | 0.93 | 0.41 | 0.73 | 0.64 | **0.10** | 0.64 | 0.54 | 0.61 | 0.91 |
| VLNP | 0.94 | 0.92 | 0.35 | 0.34 | 0.39 | 0.51 | 0.45 | 0.34 | **0.12** | 0.49 | 0.75 | 0.87 |
| PENP | 0.89 | 0.80 | 0.95 | 0.96 | 0.82 | 0.89 | 0.84 | 0.55 | 0.79 | **0.07** | 0.50 | 0.93 |
| GNG | 0.97 | 0.88 | 0.96 | 0.91 | 0.84 | 0.96 | 0.85 | 0.58 | 0.82 | 0.38 | **0.14** | 0.93 |
| OL | 0.99 | 1.00 | 0.93 | 0.98 | 0.97 | 0.98 | 0.96 | 0.91 | 0.74 | 0.99 | 0.98 | **0.37** |

model's training vocabulary (Spearman $\rho = 0.89$, $n = 12$), from 412 types (CX) to 4,273 (VLNP), and not the home MRR ($\rho = 0.07$). The target-region effect falls with the target's home MRR ($\rho = -0.81$), which is largely the ratio's denominator, and is unrelated to the target's baseline MRR ($\rho = -0.18$).

**Pair-level associations.** Across the 132 cells, the ratio correlates with the visitor's masked out-of-vocabulary share ($\rho = -0.54$), with the share of target neurons whose type the visitor knows ($\rho = 0.56$), with the share of shared F2 neurons ($\rho = 0.57$) and with type-vocabulary Jaccard similarity ($\rho = 0.55$); all permutation $p \leq 0.001$ (2,000 permutations of region labels), and $|\rho|$ 0.48–0.56 after removing row and column effects. These predictors correlate with each other at 0.84–0.98; partial correlations of shared neurons given type coverage (0.086) and of coverage given shared neurons (0.097) are near zero, so the two cannot be separated. With target fixed effects, adding the masked out-of-vocabulary share raises $R^2$ from 0.39 to 0.70 (slope −0.39). Extrapolating each column to its home model's out-of-vocabulary share predicts a mean ratio of 0.81, not 1; only four visitor cells have an out-of-vocabulary share at or below 0.30, so this extrapolation is weak.

**Asymmetry.** Transfer from $X$ to $Y$ and from $Y$ to $X$ are weakly related ($\rho = 0.18$ over 66 pairs; mean absolute difference 0.14). The difference tracks the difference in the two regions' home MRR ($\rho = -0.49$).

**Transmitter composition (post hoc).** Similarity of predicted transmitter composition between regions (one minus Jensen–Shannon distance, by neuron or by synapse, in F1 or F2) is associated with transfer at $\rho = 0.13$–$0.28$ (permutation $p$ 0.07–0.51), at 0.21–0.34 after removing row and column effects, and at 0.06–0.18 ($p$ 0.10–0.56) after also controlling for type coverage. With position features removed, transfer and transmitter similarity are unrelated ($\rho = 0.04$).

#### S8 Exploratory readout on the male CNS (F3)

The regional models (seeds 0–2) were scored on 15 F3 instances per region with no baseline and no selection. F3 graphs use thresholds of 7–11 synapses (Table S1), so their difficulty is not comparable with F2's. The project report states that F3 MRR exceeded F2 development MRR in all twelve regions. Table S13 lists the five regions the report tabulates.

**Table S13:** F3 and F2 development model MRR for the five regions tabulated in the F3 report (three seeds). These values were taken from the project report; the F3 scored rows were not available to recompute them.

| Region | F3 | F2 development |
|---|---|---|
| AL | .642 | .583 |
| LH | .560 | .387 |
| OL | .700 | .569 |
| SNP | .624 | .521 |
| VLNP | .734 | .577 |

**Table S14:** F3 MRR by the male CNS's sex-specificity annotation and by whether the masked neuron's type was in the model's training vocabulary. Random and spatial masks pooled over all twelve regions. MRR is an unweighted mean of per-type MRR; "Types" counts region–type pairs; "Positives" sums scored edges over the three seeds (each edge counted three times). Intervals are BCa bootstraps over types.

| Annotation | Trained | Types | Positives | MRR | 95% interval |
|---|---|---|---|---|---|
| Isomorphic | yes | 14,100 | 8,588,310 | .724 | [.721, .727] |
| Sexually dimorphic | yes | 218 | 70,893 | .637 | [.614, .659] |
| Potentially sexually dimorphic | yes | 108 | 24,081 | .666 | [.628, .703] |
| Isomorphic | no | 3,001 | 405,516 | .502 | [.493, .511] |
| Male-specific | no | 843 | 152,169 | .474 | [.460, .489] |
| Potentially male-specific | no | 94 | 13,986 | .450 | [.409, .497] |
| Sexually dimorphic | no | 99 | 28,926 | .436 | [.394, .481] |
| Potentially sexually dimorphic | no | 37 | 2,910 | .473 | [.404, .548] |

#### S9 Attempts to merge the regional models and to route between them

Two bodies of work tried to combine the twelve regional models into one system that could score any pair of F1-trained neurons, including pairs whose synapses no single region contains ("boundary" edges). The first was exploratory and ran only on F1. The second was pre-registered and was tested on F2. Table S15 summarizes their status; neither produced a whole-brain model that matched the regional models inside regions. This work is not part of the main text's claims; it is reported here so that the record of registered analyses and confirmation reads is complete.

**Table S15:** Status of the merging and routing attempts.

| Attempt | Registration | Outcome |
|---|---|---|
| Score fusion of two regional models (pilot) | Exploratory, F1 only | Completed. Gains of +0.004 to +0.007 F1 validation MRR over one home seed, smaller than adding a second home seed; at most +0.003 over the home three-seed ensemble. |
| Within-region fusion and routing | Exploratory, F1 only | Completed. No router or mixture beat the home model's three-seed mean. |
| Innervation-profile router on boundary edges | Exploratory, F1 only | Completed on a draft F1 suite; highest boundary MRR among methods tried there. Selected on the same masks it was reported on; never scored on F2. Code not merged into the project's main code base. |
| Fixed mixture B0 | Registered comparator | Completed on F1 validation, F2 development and F2 confirmation as an opponent; not itself gated. |
| Learned router B1 | Registered candidate | Not built: its admission rule was not met. |
| Shared union encoder A | Registered system | Completed and failed: 17 of 17 boundary comparisons passed, 0 of 12 regional guards passed. |
| Retraining screen of A | Post-failure, exploratory | Completed on F2 development; no variant came close to the regional guard. |

#### S9.1 Exploratory fusion and routing on F1

These experiments are recorded in the project's research notes (the fusion pilot and the router experiments) and in an unmerged code contribution closed on 4 November 2026, which its authors described as an experiment to inform the whole-brain phase rather than production code. Its raw result files were not kept, so every number below is taken from those records and its written report, not recomputed.

**Methods.** *Score fusion.* Each model's scores were z-scored within each ranking row and combined as $(1 - w)\, z_{\text{home}} + w\, z_{\text{other}}$, with $w$ chosen leaving one instance out. A visiting model read the target region through re-encoded features with uniform innervation. *Within-region combinations* compared equal mixes, fitted fixed weights, a fitted router and an innervation-profile router against the home model's three-seed mean. *Boundary routing* used nine frozen regional models (AL, CX, GNG, INP, LH, MB, PENP, SNP, VLNP). They scored a union graph of eleven regions (48,586 neurons, 1,232,549 edges) on a draft boundary suite with 38,907 single-masked boundary edges and 100 decoys each. The profile router weighted each model by the source's and target's innervation in that model's region, normalized over models, and had no fitted parameters. Variants added a learned residual (a softmax over the log profile prior plus a linear term) or a learned gating network trained in five folds on F1 training masks. Two of the nine models were snapshots of interrupted runs (GNG at epoch 20, MB at epoch 180).

**Results.** In the two-model pilot, fusing a visiting model into a home model raised F1 validation MRR by 0.007 for the periesophageal neuropils (0.7248 to 0.7317) and 0.004 for the ventromedial neuropils (0.7187 to 0.7227). A second seed of the home model gave more (+0.011 for the ventromedial neuropils). In nine of ten regions tried, the best single visitor added 0.001 to 0.012 to a single home seed, and fitting a weight per visitor overfit. Added to the home three-seed ensemble, a visitor changed MRR by −0.0005 to +0.0029 (for example 0.7331 to 0.7348 in the ventromedial neuropils). Within regions, no router or mixture exceeded the home three-seed mean (Table S18). On boundary edges the profile router reached 0.671, against 0.541 for the best single model and 0.568 for an equal mix (Table S16). It gained 0.136 on 400 masked neurons that no model had trained on, and 50

**Table S16:** Exploratory F1 boundary results (draft eleven-region union, four mask instances, 38,907 single-masked boundary edges). Comparison models were trained on F1 with fixed settings and are not the registered baselines.

| Method | MRR |
|---|---|
| Profile router, no fitted parameters | .6705 |
| Profile router with learned residual | .6721 |
| Learned gating network (five folds) | .6602 |
| Equal-weight mix of nine models | .5678 |
| Best single regional model (VLNP) | .5414 |
| Gradient-boosted trees on region profiles | .4826 |
| Profile router, each model restricted to its own region's edges | .4598 |
| Logistic regression | .4028 |
| Configuration model | .0273 |

**Table S17:** Exploratory F1 boundary MRR by what each regional model sees (same suite as Table S16).

| Graph and profiles | Profile router | Learned gating | Equal mix |
|---|---|---|---|
| Union graph, real profiles | .6705 | .6602 | .5678 |
| Own-region edges, real profiles | .4598 | .4887 | .4216 |
| Own-region edges, uniform profiles | .3886 | .4047 | .3866 |

random reassignments of models to regions scored 0.35–0.64. Restricting each model to its own region's edges cut the router to 0.460 (Table S17). Most of the boundary gain therefore came from letting regional models read union-graph context. Setting masked neurons' innervation to uniform in both models and router lowered the router from 0.6705 to 0.6037.

**Caveats recorded with the results.** The routing rule was chosen on the same four F1 masks it was reported on. Some models had trained on neurons in the target masks (the evaluation is transductive). The learned gates trained on a holdout that left 21% of union edges. The bootstrap intervals in those records ignore dependence between rows. None of these methods was scored on F2.

#### S9.2 Pre-registered whole-brain attempt

**Union graph and boundary suites.** The union graph sums raw synapse counts over all twelve regions' labels and thresholds once (5 in F1; 3 in F2). In F1 it has 133,910 neurons and 2,699,754 edges, of which 110,373 are boundary edges, meaning union edges present in no region's own graph. In F2 it has 131,030 neurons and 2,208,985 edges (455,366 boundary edges); these F2 counts are taken from the project's records, not recomputed here. Boundary suites hide every union edge of each masked neuron and score single-masked boundary edges against 100 union decoys. They have 5 F1 validation instances (43,265 scored edges), 12 F2 development instances (191,231) and 16 F2 confirmation instances (133,514). The F2 suites skip typed instances, which could not be drawn disjointly from the regional confirmation masks. Every scorer read masked neurons' innervation as uniform. Boundary F2 suites excluded every neuron masked by, and every type drawn by, any regional confirmation suite.

**Table S18:** Exploratory within-region combinations on F1 validation (four non-typed instances).

| Method | CX | INP | VLNP |
|---|---|---|---|
| Best single seed | .7293 | .7529 | .8129 |
| Home three-seed mean | .7445 | .7654 | .8181 |
| Equal mix of five models | .7059 | .7264 | .7728 |
| Fitted fixed weights | .7422 | .7610 | .8135 |
| Fitted router | .7402 | .7609 | .8133 |
| Profile router, query profile | .7432 | .7476 | .8097 |
| Profile router, pair profile | .7432 | .7467 | .8126 |

**Systems.** *B0* is a fixed mixture. Each region's three seeds are z-scored and averaged per row, and regions are mixed with weights from $\{0, 0.25, 0.5, 1, 2, 4\}$ fitted by coordinate ascent on F1 validation and then frozen. For regional queries B0 hands the query to that region's own models, so it matches them inside regions by construction. The *seed control* is the same mixture over five seeds per region. *B1*, a learned router, was to be built only if B0, with weights fitted leaving one instance out, trailed the best single regional model or the seed control by more than B0's spread across seeds. *A* is one shared encoder trained on the F1 union graph with uniform masked innervation (200 epochs, hidden width 128), holding out the F1 validation boundary neurons. A was admitted for testing on feasibility. Its noise floor $\delta_A = 0.01$ came from five full realizations on the F1 validation boundary suite. A second team member approved it after the calibration record was frozen and could not independently recompute the spread.

**F1 validation and the router decision.** On F1 validation boundary edges, B0 with leave-one-out weights scored 0.7626 and the seed control 0.7634. The gap (+0.0008) was smaller than B0's seed spread (0.0135), so B1 was not admitted and no learned router was ever fitted. The same rule with the earlier, innervation-reading models gave the same verdict (0.7829 against 0.7840; spread 0.0138). The admission rule compared B0 only with single regional models and the seed control. It did not consider the profile router, which scored 0.8566 on the same rows (0.8063 with masked innervation uniform in the router). A scored 0.8153 there. These F1 numbers are transductive: the models trained on F1.

**F2 development.** On F2 development boundary edges, A scored 0.699, B0 0.398, the seed control 0.408 and the best single regional model (OL) 0.490. Inside regions, A fell below the regional models of the main text (Table S7, column A).

**Gate.** The integrated claim was the conjunction of a regional non-inferiority guard in each of the twelve regions and the decision rule against each boundary opponent. The guard required the BCa lower bound of mean $\log_2(\mathrm{MRR}_A/\mathrm{MRR}_{\mathrm{reference}})$ to be at least $-\delta_r$, with no sign test. It was read on 15 November 2026 and failed (Table S19). On F2 confirmation boundary edges, A (MRR 0.724) beat all 17 opponents on all 16 instances ($p = 1.5 \times 10^{-5}$ each). These were B0 (0.401), the seed control (0.413), logistic regression (0.515), XGBoost (0.456), the configuration model (0.032) and the twelve retrained regional models (0.063–0.480). Inside regions, A's interval lay wholly below zero in eleven regions, with mean MRR 0.025–0.064 below the reference. In the optic lobe the interval included zero but its lower bound fell below the margin. Three disclosures were recorded with the verdict. First, the reference models read masked innervation computed from complete reconstructions, whereas A read it as uniform. Second, the rule that adds F2 neurons missing from the union graph (243 neurons in five regions) was adopted while a requested revision was still unresolved, and its amendment text was never marked final; A's regional confirmation scores were computed after that rule was adopted. Third, the F1 validation scores that fix B0's weights were rescored and found byte-identical to those used earlier.

**Table S19:** Pre-registered whole-brain gate, regional guards (F2 confirmation, 20 instances per region). Reference: the regional models of the main text. Margin: $-\delta_r$.

| Region | Reference MRR | A MRR | Mean $\log_2$ ratio | 95% BCa | Margin |
|---|---|---|---|---|---|
| AL | .5910 | .5640 | −.1223 | [−.3925, −.0195] | −.02 |
| CX | .5956 | .5551 | −.1276 | [−.2330, −.0418] | −.03 |
| GNG | .5792 | .5359 | −.1279 | [−.2526, −.0162] | −.02 |
| INP | .5805 | .5386 | −.1517 | [−.3439, −.0521] | −.01 |
| LH | .3634 | .3350 | −.1704 | [−.2822, −.0366] | −.01 |
| LX | .5139 | .4891 | −.1327 | [−.3788, −.0097] | −.02 |
| MB | .4417 | .4164 | −.1631 | [−.2886, −.0642] | −.03 |
| OL | .5869 | .5991 | +.0313 | [−.1632, +.1281] | .00 |
| PENP | .5320 | .4808 | −.1973 | [−.4423, −.0826] | −.02 |
| SNP | .4878 | .4441 | −.1751 | [−.4297, −.0889] | −.01 |
| VLNP | .5800 | .5420 | −.1440 | [−.3067, −.0401] | .00 |
| VMNP | .5419 | .4778 | −.2481 | [−.4813, −.1356] | −.01 |

**After the failure.** A post-gate diagnosis on F2 development found that mismatched innervation inputs do not explain A's regional deficit. The retrained regional models nearly match the originals (Table S7). Scoring regional models on the union graph instead of their own region graph lowered their MRR by 0.05 to 0.17 in the seven regions where this could be computed, a change confounded with re-encoding of features. A three-variant retraining screen of A ran on F2 development, with one seed each and no confirmation read. The variants were 400 epochs, hidden width 256, and half of training decoys drawn within regions. They moved A's mean regional $\log_2$ ratio from −0.271 to between −0.234 and −0.295. Each passed 1 of 12 regions against a continuation threshold of 8, and the screen stopped. The code change for the third variant was not committed.

**Summary.** No attempt produced a whole-brain model that matched the regional models inside regions while also scoring boundary edges. A shared encoder scores boundary edges far better than mixtures of regional models, but loses 15–19% of regional MRR on F2 development (geometric mean over regions; from almost nothing in the optic lobe to about 30% in the lateral horn). Fixed mixtures preserve regional performance by construction but score boundary edges poorly. The one method that did well on boundary edges in exploratory work, the innervation-profile router, was never tested on F2.

### S10 Pre-registration record

The analysis plan has a dated amendment log. The rows that bear on the regional comparison, all made before any confirmation suite was read, are:

- 29 August 2026: rewrite to the cross-individual design; opponent tuned and chosen on F2 development.

- 12 September 2026: typed masks drawn without replacement across development and confirmation.
- 16 September 2026: decoys keep the masked endpoint.
- 24 September 2026: the rule uses single-masked edges only; baseline training rows keep 10 of 100 decoys.
- 11 October 2026: F2 is BANC v888; selection only on F1 validation and F2 development; voiding applies per region.
- 12 October 2026: per-region $\delta$ records approved by a second member; verdicts reported per region.
- 17 October 2026: baseline fit views pinned; F1 validation check and exit rules; F2 typing by match then name.
- 19 October 2026: optic-lobe lamina types never masked.
- 3 November 2026: F1 validation screens before the F2 development digest are not counted as configurations. This was adopted without a recorded review.
- 5 November 2026: F2 typed masks limited to trained types; suites rebuilt.

The confirmation score log records every read: twelve regional passes between 7 and 13 November 2026, and the whole-brain failure on 15 November 2026. Deviations and process issues found while preparing this paper are listed in the main text (main text, Table 4). They include the innervation and position inputs of the regional models that passed, computed from complete reconstructions, the order of $\delta$ approvals for SNP and VMNP, two calibration changes committed before approval, and F2 scoring of two regions before the central complex verdict was logged. They also include registered exploratory analyses that were not run.
